# Aging disrupts the coordination between mRNA and protein expression in mouse and human midbrain

**DOI:** 10.1101/2024.06.01.596950

**Authors:** Silas A. Buck, Samuel J. Mabry, Jill R. Glausier, Tabitha Banks-Tibbs, Caroline Ward, Jenesis Gayden Kozel, Chen Fu, Kenneth N. Fish, David A. Lewis, Ryan W. Logan, Zachary Freyberg

## Abstract

Age-related dopamine (DA) neuron loss is a primary feature of Parkinson’s disease. However, it remains unclear whether similar biological processes occur during healthy aging, albeit to a lesser degree. We therefore determined whether midbrain DA neurons degenerate during aging in mice and humans. In mice, we identified no changes in midbrain neuron numbers throughout aging. Despite this, we found age-related decreases in midbrain mRNA expression of tyrosine hydroxylase (*Th*), the rate limiting enzyme of DA synthesis. Among midbrain glutamatergic cells, we similarly identified age-related declines in vesicular glutamate transporter 2 (*Vglut2*) mRNA expression. In co-transmitting *Th*^+^/*Vglut2*^+^ neurons, *Th* and *Vglut2* transcripts decreased with aging. Importantly, striatal Th and Vglut2 protein expression remained unchanged. In translating our findings to humans, we found no midbrain neurodegeneration during aging and identified age-related decreases in *TH* and *VGLUT2* mRNA expression similar to mouse. Unlike mice, we discovered diminished density of striatal TH^+^ dopaminergic terminals in aged human subjects. However, TH and VGLUT2 protein expression were unchanged in the remaining striatal boutons. Finally, in contrast to *Th* and *Vglut2* mRNA, expression of most ribosomal genes in *Th*^+^ neurons was either maintained or even upregulated during aging. This suggests a homeostatic mechanism where age-related declines in transcriptional efficiency are overcome by ongoing ribosomal translation. Overall, we demonstrate species-conserved transcriptional effects of aging in midbrain dopaminergic and glutamatergic neurons that are not accompanied by marked cell death or lower striatal protein expression. This opens the door to novel therapeutic approaches to maintain neurotransmission and bolster neuronal resilience.

## Introduction

Aging is the greatest risk factor in several prevalent neurodegenerative diseases, including Parkinson’s Disease (PD)^1, 2^. Yet, even in healthy aging, previous work has shown that significant changes occur in brain neurotransmission, including within the dopamine (DA) system^3^. Indeed, there is a decrease in the synthesis^4^ and transport^5^ of DA, as well as in DA receptor expression^6^ across aging in humans. Furthermore, such changes appear to be conserved across species with reports of DA neuron loss during healthy aging in *Drosophila*^7^ and in mice^8^. Nevertheless, the exact relationships between loss of DA function and potential DA neuron degeneration remain a subject of active debate, particularly in humans^9^. Clearly, the pathologies underlying age-related neurodegenerative disorders are more severe than the changes in the DA system associated with healthy aging. Yet many key questions remain concerning the links between the aging process and DA neuron health.

Previous attempts at examining markers of DA neuron function or identity throughout the course of aging have often yielded conflicting results. Though some work showed aging-related decreases in midbrain DA neuron markers^10–17^, other studies demonstrated no difference^18–22^. Nevertheless, there is an emerging consensus that certain aspects of dopaminergic cell biology are modified by the aging process. Besides neuromelanin, other DA neuron markers have also been studied in aging. There is age-related loss of DA transporter (DAT) mRNA expression, suggesting potential impairments in DA neuron function and/or loss^13, 14^. Decreased expression of tyrosine hydroxylase (TH), the rate limiting enzyme in DA biosynthesis, has also been reported during aging^11^. However, loss or diminishment of TH expression does not necessarily represent frank DA neurodegeneration since cells with little TH expression can still express other markers of DA biosynthesis like aromatic L-amino acid decarboxylase (AADC) or vesicular storage such as vesicular monoamine transporter 2 (VMAT2)^23–25^. Moreover, despite the consensus that mRNA and protein expression are closely coordinated, mRNA and protein expression can diverge, via changes in trafficking, translation, and mRNA/protein degradation^26^. To date, it remains unclear if or how aging alters such coordination between mRNA and protein expression within the DA system. Moreover, are there neuroprotective factors that preserve DA neuron survival and function during aging?

Important clues come from models of DA neuron aging in *Drosophila*. We showed that the *Drosophila* ortholog of the vesicular glutamate transporter 2 (VGLUT2), dVGLUT, protected DA neurons from age-related neurodegeneration^27^. This raises the question of VGLUT’s generalizability as a protective factor for DA neurons in aging. While studies have investigated VGLUT2’s roles in mammalian PD models, it remains unknown whether VGLUT2-expressing midbrain neurons, including TH^+^/VGLUT2^+^ DA neurons and purely glutamatergic neurons (*i.e*., TH^-^/VGLUT2^+^), are more resilient to the effects of normal aging in rodent or human brain.

Here, we investigated the relationships between age-related differences in midbrain DA neuron *TH* gene expression and cell survival in both mouse and human brain across normal aging. We employed multiplex RNAscope fluorescent *in situ* hybridization which quantitatively measures mRNA expression at single-cell resolution across the midbrain. In parallel, we examined effects of aging on *VGLUT2* expression and survival of midbrain glutamatergic neurons, including subpopulations of TH^+^/VGLUT2^+^ DA neurons as well as purely glutamatergic TH^-^/VGLUT2^+^ cells. In mice, midbrain *Th* and *Vglut2* mRNA expression significantly decreased across aging while total cell density remained unchanged. This suggests that age-related decreases in *Th* and *Vglut2* mRNA expression are not a consequence of neuronal loss in mouse midbrain. By contrast, Th and Vglut2 protein levels in mouse striatum did not significantly change as a function of age. We discovered that, like mice, human midbrains exhibited age-associated declines in *TH* and *VGLUT2* mRNA expression. Yet, unlike mice, aged human subjects exhibited decreased density of striatal TH^+^ dopaminergic terminals, while striatal VGLUT2^+^ terminal density remained unaffected. Importantly, in spite of this age-related decrease in dopaminergic terminal density, levels of TH and VGLUT2 protein expression did not decline in the remaining striatal terminals of aged individuals. These results point to a conserved mechanism in aging mice and humans by which protein expression is preserved despite dwindling mRNA expression of key components of the dopaminergic and glutamatergic neurotransmission machinery. Importantly, transcriptomic analyses revealed preservation and even upregulation of ribosomal genes in mouse DA neurons during aging, offering a mechanism to maintain protein expression. Better understanding of these conserved mechanisms may lead to more effective strategies to boost midbrain neuron resilience throughout the aging process.

## Methods

### Sex as a biological variable

We conducted this study on midbrain and striatum of both male and female mice and humans.

### Mice

Young (3-5 months old; N=5 males, 5 females), middle-aged (10-12 months old; N=3 males, 4 females), and aged (19-21 months old; N=4 males, 4 females) wild-type C57BL/6J mice (The Jackson Laboratory, Bar Harbor, ME; JAX no. 000664) were maintained on a 12:12 hour light:dark cycle with food and water available *ad libitum*. Age designations for the respective age-groups were determined according to previously described criteria^28^. Animals were euthanized either with carbon dioxide followed by decapitation (for RNAscope fluorescent *in situ* hybridization experiments) or with isofluorane and transcardially perfused with 4% paraformaldehyde (for immunohistochemistry experiments). All mouse experiments were approved by the University of Pittsburgh Institutional Animal Care and Use Committee (Protocol# 18012220). Animals were cared for in accordance with all appropriate NIH animal care guidelines and according to the ARRIVE guidelines for reporting animal research^29–31^. All efforts were made to ameliorate animal suffering.

### Human subjects

Brain specimens (N=12 subjects) were obtained through the University of Pittsburgh Brain Tissue Donation Program following consent from next-of-kin, during autopsies conducted at the Allegheny County (Pittsburgh, PA) or the Davidson County (Nashville, TN) Office of the Medical Examiner. An independent committee of experienced research clinicians confirmed the absence of lifetime psychiatric and neurologic diagnoses for all subjects based on medical and neuropathological records, toxicology reports, as well as structured diagnostic interviews conducted with family members of the deceased^32, 33^. Subjects were categorized as young (16-21 years, N=4), middle-aged (45-49 years, N=4), and aged (65-74 years, N=4). Each age group included two male and two female subjects, and subject groups did not differ in mean postmortem interval (PMI), RNA integrity number (RIN) or brain pH (Table 1). One aged subject was removed from analysis due to outlier cell density levels of *TH* mRNA (*TH*^+^) cells and co-expressing *TH* and *VGLUT2* mRNA (*TH*^+^/*VGLUT2*^+^) cells, as determined by the Robust Regression and Outlier (ROUT) method^34^, with a Q coefficient of 1 (Supplementary Figure 1). All analyses were therefore performed on the 11 remaining subjects, and the subject groups did not differ in mean PMI, RIN, or brain pH (all F_2,8_<0.3 and all P>0.75). All procedures were approved by the University of Pittsburgh’s Committee for Oversight of Research and Clinical Training Involving Decedents and Institutional Review Board for Biomedical Research.

**Table 1.** Summary of demographic and postmortem characteristics of human subjects. Values are mean (±SD). PMI, postmortem interval; RIN, RNA integrity number; *aged female outlier subject was excluded from analyses.

| Subject Group | Age | Sex (M/F) | PMI, hr | pH | RIN |
| --- | --- | --- | --- | --- | --- |
| Young | 19.0 (2.9) | 2/2 | 17.1 (6.3) | 6.9 (0.1) | 8.2 (0.5) |
| Middle-Aged | 47.0 (1.8) | 2/2 | 14.3 (5.2) | 6.7 (0.2) | 8.1 (0.5) |
| **Aged** | 68.0 (4.1) | 2/2* | 13.2 (5.0) | 6.6 (0.2) | 8.2 (0.5) |
| **Statistics** | | | $F_{2,8} = 0.29, P = 0.76$ | $F_{2,8} = 0.27, P = 0.77$ | $F_{2,8} = 0.2, P = 0.83$ |

### RNAscope multiplex fluorescent *in situ* hybridization

#### Tissue preparation

For mouse brain samples, brains were immediately frozen after collection. This was followed by sectioning into 20 μm-thick sections at −3.2 to −3.6 mm from Bregma using a cryostat (Leica Biosystems, Wetzlar, Germany). Midbrain sections containing ventral tegmental area (VTA) and substantia nigra pars compacta (SNc) were then mounted onto Superfrost Plus slides (Fisher Scientific, Hampton, NH) and stored at −80°C until staining. For human brain samples, right hemisphere midbrain containing the VTA and substantia nigra (SN) was cut at a 20 μm thickness via cryostat, mounted onto Superfrost Plus slides and stored at −80°C until processing.

#### Multiplex RNAscope

Probes for multiplex fluorescent *in situ* hybridization were designed by Advanced Cell Diagnostics, Inc. (Hayward, CA) to detect mRNAs encoding TH (*TH* gene in humans, Catalog No. 441651-C2; *Th* in mice, Catalog No. 317621-C2), VGLUT2 (*SLC17A6* gene in humans, Catalog No. 415671; *Slc17a6* gene in mice, Catalog No. 319171), and *Ribosomal protein L6* (*Rpl6* gene in mice, Catalog No. 544371). All mouse and human tissue sections were processed using the RNAscope Assay V1 kit (Advanced Cell Diagnostics) according to the manufacturer’s protocol. One section from each subject was processed on the same day to reduce batch effects. Briefly, 2 adjacent human or mouse tissue sections were fixed for 15 min in ice-cold 4% paraformaldehyde, incubated in a protease treatment, followed by probe hybridization to their target mRNAs (2 hours, 40°C). The sections were exposed to a series of incubations that amplified the target probes, and then counterstained with DAPI for cell nucleus labeling. Human brain sections were labeled for *TH* and *VGLUT2* mRNAs, while mouse brain sections were labeled for *Th*, *Vglut2*, and *Rpl6* mRNAs. *VGLUT2/Vglut2*, *TH/Th*, and *Rpl6* mRNAs were subsequently detected with Alexa 488, Atto 550, and Alexa 647 dyes, respectively.

### Fluorescence microscopy of mRNA expression

RNAscope images of human and mouse brain sections were acquired with an Olympus IX81 inverted fluorescence microscope using a 60× 1.4 NA SC oil immersion objective. The microscope was additionally equipped with a Hamamatsu ORCA-Flash4.0 CCD camera (Hamamatsu Corporation, Bridgewater, NJ) and high-precision BioPrecision2 XYZ motorized stage with linear XYZ encoders (Ludl Electronic Products Ltd, Hawthorne, NJ). The VTA and SNc of mice were identified using stereotaxic coordinates and well-established anatomic landmarks^35^. For both mouse and human samples, 3D image stacks (2 adjacent sections/subject; 2048×2048 pixels; 0.2 μm z-steps) of 100% of the tissue thickness were taken spanning the entire medial-lateral and dorsal-ventral axes. Image sites were systematically and randomly selected using a grid of 100 μm^2^ frames spaced by 500 μm (for mice) or 350 μm (for humans). This imaging scheme resulted in the imaging of 25-40 sites/section in mice and 100-300 sites/section in humans. Both hemispheres were imaged in mouse samples while the right hemisphere was imaged in human samples. Image collection was controlled by Slidebook 6.0 (Intelligent Imaging Innovations, Inc., Denver, CO). The z-stacks were collected using optimal exposure settings (*i.e.*, those that yielded the greatest dynamic range with no saturated pixels), with differences in exposures normalized during image processing. To manage native tissue fluorescence in our human postmortem tissue samples from lipofuscin, an intracellular lysosomal protein that accumulates with age^27, 36, 37^, we effectively excluded lipofuscin signal from our RNAscope probe-specific signals using a previously described approach^38–40^. Briefly, we imaged lipofuscin in human sections using a fourth visible channel (excitation/emission: 405 nm/647 nm) and masked the lipofuscin signal using an optimal threshold value. In studies examining midbrain *Rpl6* mRNA expression, images of whole coronal sections containing midbrain were acquired using an Olympus VS120 automated slide scanner (Olympus, Center Valley, PA) equipped with a 20× 0.8 NA dry objective. 3D images were composed of z-stacks composed of 3 z-planes spaced 1 μm apart. Identical exposure settings and magnifications were consistently applied to all slides.

### Image analysis of mRNA expression

Mouse and human brain RNAscope data imaged on the Olympus inverted fluorescence microscope were initially analyzed via Slidebook and Matlab software (MathWorks, Natick, MA). A Gaussian channel was constructed for each channel by calculating a difference of Gaussians using sigma values of 0.7 and 2. Average intensity projections of every 3D image z-stack were created by averaging intensity values within each Gaussian channel to assemble a 2D image of DAPI-stained cells alongside the respective mRNA transcripts (*TH*/*VGLUT2* for human samples and *Th*/*Vglut2* for mouse samples). Mouse brain data acquired from the Olympus V120 slide scanner were similarly converted into 2D maximum intensity projections via Olympus ASW software (version 3).

### DAPI-stained cell and mRNA quantification

To quantify DAPI-stained cell numbers and mRNA expression in both mouse and human brain tissues, 2D projection images were separated into quantitative TIFF files and analyzed via HALO image analysis software equipped with a fluorescent *in situ* hybridization module (version 3.0, Indica Labs, Albuquerque, NM). For mouse and human brain data acquired on the Olympus inverted fluorescence microscope, DAPI-labeled cell nuclei and fluorescent grains representing mRNA transcripts were quantified using the following parameters for inclusion in our counts based on the following thresholding: any object 40-500 μm^2^ for DAPI and 0.1-0.5 μm^2^ for mRNA grains. Objects from the other channels that overlapped with lipofuscin were eliminated from analyses by subtracting the lipofuscin Gaussian channel from the other channels. Histograms of mRNA grain levels for *TH* and *VGLUT2* in all human cells as well as *Th* and *Vglut2* in all mouse cells analyzed are shown in Supplementary Figure 2. We determined positive cell expression levels as previously described^41^, with some modifications. To determine the minimum number of mRNA grains for a labeled cell to be considered positive for expression, we tested different thresholds of 1.5x, 3x, and 5x the number of mRNA grains above background levels (*i.e.*, numbers of grains expressed in a typical cell volume, Supplementary Figure 3). Since there were no differences in relative *TH*^+^ and *TH*^+^/*VGLUT2*^+^ cell density changes between the thresholds (P>0.05; Supplementary Figure 3), a minimum threshold of 3x the background expression level was selected as the threshold for quantifying positive cells. All mRNA grains within 5 μm of the DAPI-stained nucleus edge were considered as belonging to the respective cell. Wherever necessary, this 5 μm border was reduced to prevent overlap between neighboring cells during analysis. RNAscope imaging data acquired on the Olympus VS120 slide scanner were similarly analyzed as above. Briefly, nuclei were quantified as DAPI-stained objects with a minimum cytoplasmic radius set at 5 μm. Puncta corresponding to each respective mRNA probe were identified as any 0.03-0.15 μm^2^ object. We used custom code to define cells that were positively labeled by the respective mRNA probes. We determined that a minimum threshold of 2x above baseline provided optimal signal-to-noise in our data, enabling accurate detection of labeled cells. Cell densities were calculated by dividing the number of positive cells by the total area (in mm^2^ units). In testing for potential sex effects, there were no main effects of sex on *TH* expression (all P>0.05), thus males and females were combined for further analyses.

### Mouse and Human Tissue

#### Mice

Mice were perfused with 4% paraformaldehyde in PBS, with extracted brains subsequently post-fixed by immersion in 4% paraformaldehyde in PBS for 24 hours. Brains were then transferred to 30% sucrose and stored at 4°C for at least 48 hours before sectioning. Brains were sectioned (35 μm thickness, Anterior-Posterior: 1.6-1.2 mm from Bregma, 2 sections/animal separated by 0.2 mm) using a cryostat and maintained at −20°C in ethylene glycol cryoprotectant prior to immunohistochemistry.

#### Humans

The left hemisphere of each brain was blocked coronally, fixed in cold 4% paraformaldehyde for 48 hours, immersed in a series of graded sucrose solutions and then cryoprotected. Human tissue blocks containing the striatum were sectioned coronally (40 μm thickness) using a cryostat and stored in antifreeze solution at −30°C until processing.

### Immunohistochemistry

#### Immunohistochemical labeling

Free-floating striatal mouse sections were washed with PBS and blocked in PBS supplemented with 10% Normal Donkey Serum (NDS) and 1% Triton X-100 (1 hour, room temperature). Free-floating human striatal sections were washed with PBS and incubated in phosphate buffer with 1% sodium borohydride (30 min, room temperature). Sections were then incubated in PBS with 0.3% Triton X-100 for 30 min and blocked in PBS supplemented with 20% NDS (2 hours, room temperature). Both mouse and human brain sections were labeled for TH (mouse monoclonal, clone LNC1, 1:2000, Catalog No. MAB318, EMD Millipore, Burlington, MA) and VGLUT2 (rabbit polyclonal, 1:500, Catalog No. 135403, Synaptic Systems, Göttingen, Germany). The respective specificities of anti-TH and anti-VGLUT2 primary antibodies have been previously described^42, 43^. Samples were incubated with primary antibodies for either 24 hours (for mouse samples) or 72 hours (for human samples) at 4°C. The respective mouse and human brain sections were subsequently incubated with Invitrogen donkey anti-rabbit Alexa Fluor 488 (Catalog No. A-21206; Invitrogen, Carlsbad, CA) for VGLUT2, and donkey anti-mouse Alexa Fluor 555 for TH (Catalog No. A-31570; Invitrogen) secondary antibodies. Samples were incubated with secondary antibodies for either 2 hours (mouse sections) or 24 hours (human sections). Nucleus accumbens and caudate/putamen of each sample were labeled together and, for human subjects, subjects within a triad were processed together. Labeled sections were then washed in PBS and mounted onto glass slides for imaging.

### Confocal microscopy and image analysis of protein expression

The number of sampling sites in human striatum was chosen on the basis of pilot immunohistochemistry in mouse striatum comparing TH and VGLUT2 protein co-localization in the dorsal striatum (caudate/putamen) versus the ventral striatum (nucleus accumbens). 8-12 sites were imaged (covering the entirety of each region, 2 sections per animal) in both the dorsal and ventral striatum for young, middle-aged, and aged mice (N = 8-10 per group). Bayesian analyses of TH and VGLUT2 puncta density, intensity, and co-localization demonstrated substantial evidence of absence (Table 2). Additionally, a running means analysis of mice showed the means of the dependent measures converged before 8 sites had been imaged (Supplementary Figure 4). Together, these results in mice guided our decision to similarly sample 8-12 sites per region in human striatum. Consequently, we employed a sampling grid covering the entire region of interest which was composed of 100 μm × 100 μm sampling boxes separated by 2000 μm to achieve the desired sampling of 8-12 sites per section (Supplementary Figure 5).

**Table 2.** Bayesian analyses of TH and VGLUT2 puncta density, intensity, and co-localization. Bayesian 2-way ANOVA (using age and brain region as main effects) in mice demonstrating, with the exception of VGLUT2 puncta density, substantial (BF_01_ = 3.2-10) or strong (BF_01_ = 10-100) evidence of absence.

|  | TH | VGLUT2 |
| --- | --- | --- |
| **Puncta Density** | $\text{BF}_{01} = 14.317$ | $\text{BF}_{01} = 2.948$ |
| **Puncta Intensity** | $\text{BF}_{01} = 9.482$ | $\text{BF}_{01} = 15.091$ |
| **TH Co-localization with VGLUT2** | $\text{BF}_{01} = 70.334$ | |

Immunohistochemical labeling images were collected on an Olympus IX81 inverted microscope equipped with a 60× 1.40 NA SC oil immersion objective, an Olympus spinning disk confocal unit, Hamamatsu ORCA-Flash4.0 CCD camera, and high precision BioPrecision2 XYZ motorized stage with linear XYZ encoders; equipment was controlled by SlideBook 6.0 software. 3D image stacks (1024 × 1024 pixels, 0.25 μm z-steps) were acquired over the middle 50% of the total thickness of the tissue sections. Caudate/putamen and nucleus accumbens in mouse or human striatum were identified using previously described stereotactic coordinates^35^ as well as according to anatomic features. Image stacks were randomly selected using a sampling grid of 100 μm^2^ frames spaced by 750 μm (for mice) or 2000 μm (for humans), leading to imaging of 8-12 sites per section as detailed above. An identical exposure time was maintained across all images and sections. Each fluorescent channel was deconvolved using Autoquant’s Blind Deconvolution algorithm. Then, a Gaussian channel was made for each deconvolved channel by calculating a difference of Gaussians using sigma values of 0.7 and 2. The Gaussian channel was used for data segmentation using Matlab. For segmentation, object masks were created using a size-gating of 0.05-0.7μm^3^ as previously^40^. For human sections, lipofuscin was imaged and subtracted from other channels as described above. The mean density of TH object masks and VGLUT2 object masks was calculated for every sample in each striatal region. TH and VGLUT2 fluorescence intensities, reflecting the relative amount of protein^44^, were extracted from their respective object masks to calculate mean intensity for each subject and striatal region. The density of TH puncta that overlapped with VGLUT2 puncta by at least 20% was analyzed across age groups. As with RNAscope imaging, overlapping threshold did not have an impact on the relative differences in overlapping puncta between age groups and brain region (Supplementary Figure 6). TH/VGLUT2 overlapping masks were included in analyses of TH object masks and VGLUT2 object masks.

### Transcriptomic analyses

We conducted transcriptomic analyses using a previously deposited dataset of unbiased high-throughput single-cell RNA sequencing (scRNA-seq) data from brains of young versus aged wild-type male C57BL/6J mice (GSE129788)^45^. This dataset consisted of young (2-3 months old; N=8 males) and aged mice (21-22 months old; N=8 males); the ages of the mice constituting these age groups overlapped with the mice used in our RNAscope/IHC analyses. In total, we detected 797 Th^+^ neurons (TPM_Th_>0). 169 known ribosomal genes were expressed in at least one group of either young or aged animals. We filtered out genes expressed in <10 cells per age group, resulting in a final set of 141 ribosomal genes for analysis. The distribution of transcripts per million (TPMs) in both young and aged groups was represented by violin plots which were generated by R package ggplot2^46^. Ratios of averaged expression for all analyzed ribosomal genes in aged versus young groups were graphed as bar plots.

### Statistical analyses

Immunohistochemistry and RNAscope data were analyzed using a two-way analysis of variance (ANOVA) to test for both main effects of and interaction between age group and brain region. Bonferroni post hoc analyses were performed to investigate significant effects. A three-way ANOVA was conducted to test for main effects of *Rpl6* mRNA expression, age groups, and cell types. In mice, since we found no significant differences between males and females across all examined age groups (p>0.05), sex was not included as a main effect in subsequent analyses. For RNAscope and immunohistochemistry of postmortem human brain samples, two-way analysis of covariance (ANCOVA) models were performed with age group and brain region as main effects and PMI, pH and RIN as covariates. For all ANCOVAs, reported statistics only included the covariates that showed statistical significance; as such, the reported degrees of freedom varied across analyses. Correlations between age and mRNA expression of *TH* and *VGLUT2* in the VTA and SNc of human subjects were also analyzed. Bayesian analyses of TH^+^ and VGLUT2^+^ protein puncta density, intensity, and co-localization were performed to evaluate evidence of absence. Additionally, running means analyses were used to determine an adequate number of imaging sites in immunohistochemically labeled brain tissue. For all statistics, significance was defined as P<0.05. All statistical analyses were carried out using GraphPad Prism software (Version 8.2) and SPSS (version 26; IBM, Armonk, NY).

## Results

### *Th* mRNA expression decreases across aging in mouse midbrain

We first examined the impact of healthy aging on the density of midbrain DA neurons expressing *Th* mRNA (*i.e*., *Th*^+^ cells) via RNAscope in young, middle-aged, and aged wild-type mice (Figure 1A-H). Aging significantly impacted the density of *Th*^+^ cells, with a 48% decreased density of *Th*^+^ cells in aged mice compared to young mice (Figure 1G). Interestingly, there were no significant differences between midbrain regions, with the VTA and SNc exhibiting similarly diminished *Th*^+^ neuron densities (Figure 1G). Furthermore, we discovered age-related changes in *Th* mRNA expression, with aged animals having diminished *Th* expression (Figure 1H). Post hoc analyses revealed a progressive age-related decline in *Th* mRNA expression where the aged group exhibited 9% fewer *Th* mRNA grains per cell compared to young and 15% fewer grains versus middle-aged groups.

**Figure 1.**
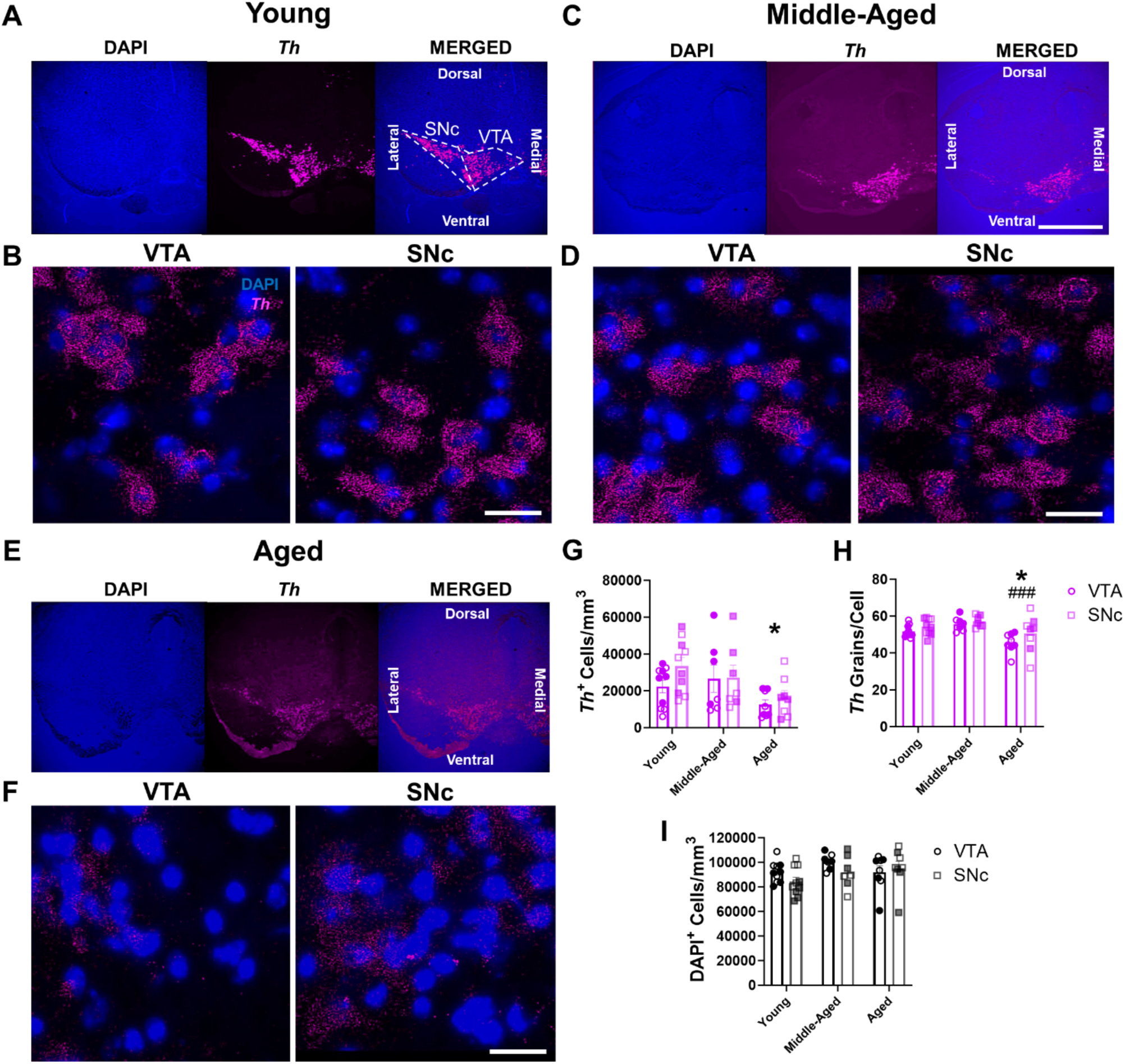
Age-related decreases in *Th* mRNA expression in mouse VTA and SNc. **(A-F)** Representative RNAscope images of *Th* mRNA expression (in magenta) and associated DAPI-labeled cell nuclei (in blue) in mouse midbrain across aging. 4× images of young **(A)**, middle-aged **(C)**, and aged mice **(E)**; scale bar=500μm. Enlarged 60× images show *Th* expression and nuclei within VTA and SNc midbrain subregions in young **(B)**, middle-aged **(D)**, and aged **(F)** animals; scale bar=25μm. **(G)** There were significant age-related decreases in the cell density of *Th*^+^ dopaminergic cells in the VTA and SNc (F_2,44_=4.8, P=0.012 for effect of age group; F_1,44_=1.7, P=0.2 for effect of brain region; F_2,44_=0.7, P=0.52 for effect of interaction). **(H)** There were significant age-related decreases in *Th* mRNA grain numbers per cell of *Th*^+^ DAergic cells in the VTA and SNc (F_2,44_=8.9, P=0.0006 for effect of age group; F_1,44_=3.6, P=0.06 for effect of brain region; F_2,44_=0.4, P=0.66 for effect of interaction). **(I)** There was no significant change in DAPI-labeled nucleus density with age in either the VTA or SNc (F_2,44_=2.1, P=0.14 for effect of age group; F_1,44_=1.6, P=0.21 for effect of brain region; F_2,44_=1.2, P=0.31 for effect of interaction). Shaded symbols represent female animals. Bars represent mean±SEM with points representing individual animals; N=7-10 per group. *P<0.05 compared to young age, ^###^P<0.001 compared to middle-aged, via Bonferroni post hoc test.

We next determined whether the reduced density of *Th*^+^ cells in the VTA and SNc in the aged group was the product of lower total cell density overall, or whether this reduction represented a loss of *Th* expression in surviving neurons. To test this, we measured total cell density within the midbrain via DAPI nuclear staining in the same brain sections. We found no significant differences in nucleus density either as a function of age group or region (Figure 1I). Importantly, these data demonstrated the absence of cell loss across aging in mouse midbrain. Taken together, age-related decreases in *Th* mRNA grains, coupled with no significant alterations in DAPI^+^ cell density, strongly suggest that diminished *Th*^+^ cell density across aging is due to decreased *Th* mRNA expression rather than actual dopaminergic cell death.

### Regional and cell subpopulation differences in mouse midbrain *Th* mRNA expression across aging

We investigated whether specific midbrain regions and/or neuronal subpopulations were more prone to age-related mRNA expression changes than others. In addition to dopaminergic cells that exclusively transmit DA, there are also populations of midbrain DA neurons that co-transmit glutamate and express *Vglut2*. Initially focusing on *Th*^+^ cells that do not co-express *Vglut2* (*Th*^+^/*Vglut2*^-^), we found an overall higher density of *Th*^+^/*Vglut2*^-^ neurons in the lateral VTA compared to the medial VTA (Figure 2A-C). However, there were no significant differences in the density of VTA *Th*^+^/*Vglut2*^-^ neurons between age groups (Figure 2C). In contrast, in the SNc, we found a significant effect of age on *Th*^+^/*Vglut2*^-^ neuron density. Aged animals exhibited a 51% decrease compared to young animals in cells with identifiable *Th* mRNA signal but no differences between SNc sub-regions (*i.e.*, medial versus lateral SNc) (Figure 2D). These findings show that, in addition to overall loss of *Th* mRNA signal in cells with age, there is a significant regional component to these age-related changes in *Th* mRNA expression within the ventral midbrain.

**Figure 2.**
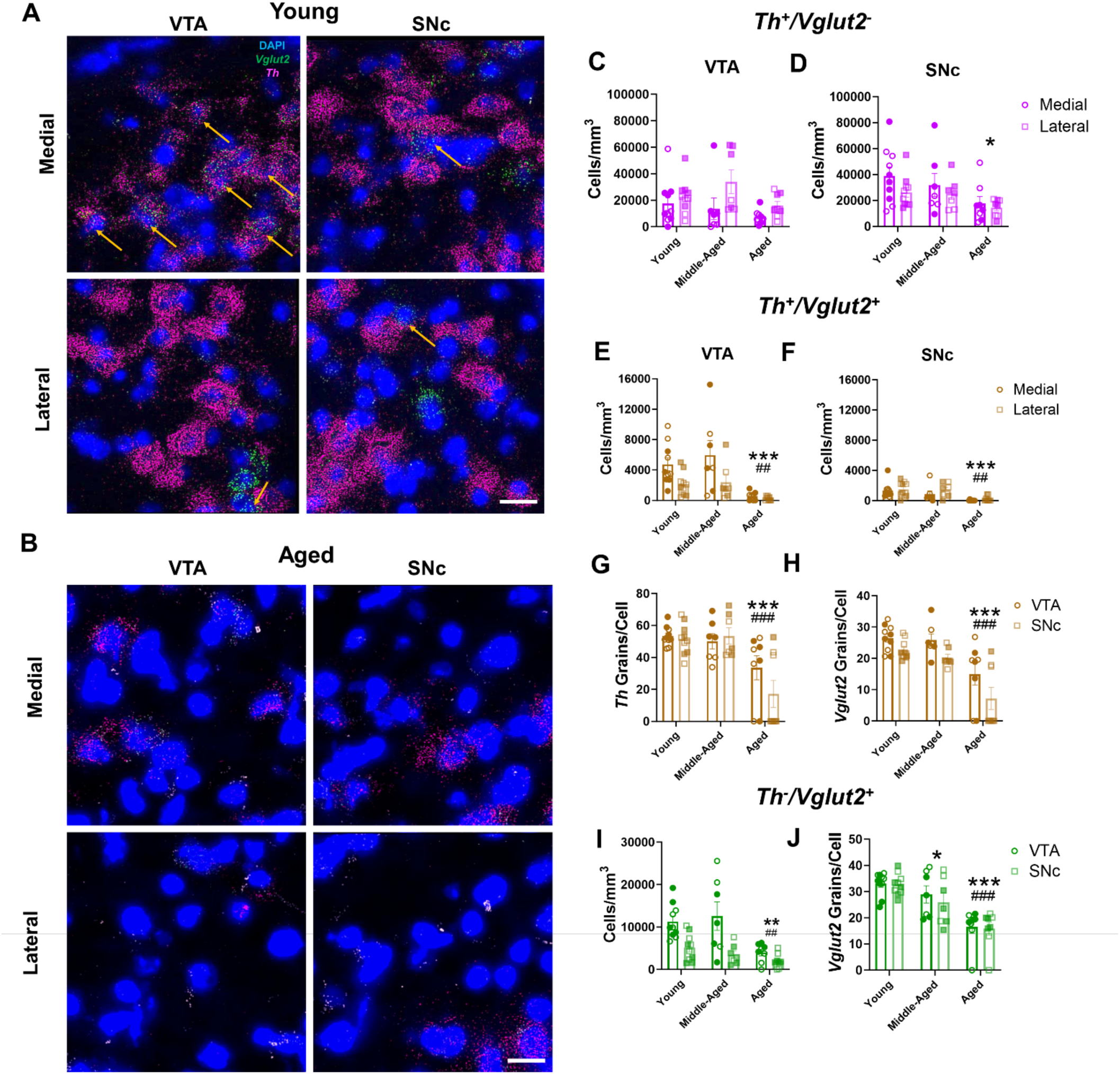
*Th* and *Vglut2* mRNA expression decrease across aging in distinct midbrain neuron subpopulations. **(A-B)** Representative 60× multiplex RNAscope images of *Th* (in magenta) and *Vglut2* (in green) mRNA expression alongside DAPI nuclear stain (in blue) in the VTA and SNc of young **(A)** and aged **(B)** wild-type mice. *Th*^+^/*Vglut*2^+^ co-expressing midbrain neurons are marked by yellow arrows; scale bar=25μm. **(C)** There were significantly more *Th^+^/Vglut2^-^* cells in the lateral VTA compared to the medial VTA (F_2,44_=2.4, P=0.11 for effect of age group; F_1,44_=6.2, P=0.017 for effect of subregion; F_2,44_=1.0, P=0.37 for effect of interaction). **(D)** There was a significant age-related decrease in the cell density of *Th*^+^ neurons in the SNc (F_2,44_=4.9, P=0.01 for effect of age; F_2,44_=2.6, P=0.11 for effect of subregion; F_2,44_=0.38, P=0.68 for effect of interaction). **(E-F)** There was an age-related decrease in the cell density of *Th*^+^/*Vglut2*^+^ neurons in the VTA (**E**: F_2,44_=9.9, P=0.0003 for effect of age; F_1,44_=9.1, P=0.004 for effect of subregion; F_2,44_=1.7, P=0.20 for effect of interaction) and SNc (**F**: F_2,44_=10.3, P=0.0002 for effect of age; F_1,44_=1.6, P=0.21 for effect of region; F_2,44_=0.2, P=0.82 for effect of interaction) of aged mice compared to young and middle-aged animals. **(G)** *Th* mRNA expression decreased within midbrain *Th*^+^/*Vglut2*^+^ cells in aged mice (F_2,44_=16.62, P<0.0001 for effect of age; F_1,44_=1.247, P=0.27 for effect of region; F_2,44_=1.736, P=0.19 for effect of interaction). **(H)** *Vglut2* mRNA grain expression decreased within midbrain *Th*^+^/*Vglut2*^+^ cells in aged mice. Additionally, *Vglut2* mRNA grain expression was greater in the VTA compared to SNc (F_2,44_=21.3, P<0.0001 for effect of age; F_1,44_=79.758, P=0.032 for effect of brain region; F_2,44_=0.40, P=0.68 for effect of interaction). **(I)** There was a lower cell density of purely glutamatergic *Th*^-^/*Vglut2*^+^ neurons in the VTA and SNc of aged mice compared to young and middle-aged mice (F_2,44_=7.9, P=0.0012 for effect of age; F_1,44_=22.1, P<0.001 for effect of region; F_2,44_=2.7, P=0.08 for effect of interaction). **(J)** There was a significant decrease in *Vglut2* mRNA grains per *Th*^-^/*Vglut2*^+^ cells in middle-aged compared to young mice. Aged mice exhibited a further decrease in *Vglut2* mRNA grains compared to young and middle-aged mice. (F_2,44_=28.12, P<0.001 for effect of age; F_1,44_=0.52, P=0.48 for effect of region; F_2,42_=0.17, P=0.84 for effect of interaction). Shaded symbols represent females and open symbols represent males. Bars represent mean±SEM with points representing individual animals; N=7-10 per group. *P<0.05, **P<0.01, ***P<0.001 compared to young age, ^#^P<0.05, ^##^P<0.01 compared to middle-aged, ^###^P<0.001 compared to middle-aged, via Bonferroni post hoc test.

### Age-related decreases in mouse *Th* and *Vglut2* mRNA expression in co-transmitting DA/glutamate neurons

We next examined age-related changes in both *Th* and *Vglut2* mRNA expression within the subpopulation of midbrain *Vglut2*^+^ DA neurons (*i.e.*, *Th^+^/Vglut2*^+^ neurons). We found more cells co-expressing *Th* and *Vglut2* mRNA in the medial VTA compared to lateral VTA (Figure 2E), consistent with earlier work^47–49^. There were also significantly fewer *Th*^+^/*Vglut2*^+^ cells in the VTA of aged mice compared to both young (88% decrease) and middle-aged (89% decrease) animals (Figure 2E). Like the VTA, there was a decrease in *Th*^+^/*Vglut2*^+^ cell density in the SNc across aging, with the aged group having fewer *Th*^+^/*Vglut2*^+^ cells compared to both young (90% decrease) and middle-aged (88% decrease) mice (Figure 2F).

Within *Th*^+^/*Vglut2*^+^ cells, aged mice possessed 48% fewer *Th* mRNA grains per cell versus young mice and 51% fewer compared to middle-aged mice (Figure 2G). For *Vglut2* mRNA grains within *Th*^+^/*Vglut2*^+^ co-expressing cells, aged mice had 45% fewer mRNA grains per cell compared to young mice and 52% fewer compared to middle-aged mice (Figure 2H). Furthermore, there were fewer *Vglut2* mRNA grains within *Th*^+^/*Vglut2*^+^ co-expressing cells in the SNc compared to the VTA (Figure 2H). Overall, though *Th^+^/Vglut2*^+^ neurons are relatively more resilient to neurodegeneration^50^, these data suggest that DA neuron *Vglut2* mRNA expression does not modify or protect against age-related decreases in *Th* mRNA expression, especially since *Vglut2* mRNA expression also decreases with age.

### *Vglut2* mRNA expression in non-dopaminergic glutamatergic mouse midbrain neurons diminishes across aging

We determined whether *Vglut2* mRNA expression decreased with age in purely glutamatergic *Vglut2*^+^ (*Th*^-^/*Vglut2*^+^) midbrain neurons. Similar to *Th*^+^/*Vglut2*^+^ cells, there was diminished density of VTA *Th*^-^/*Vglut2*^+^ neurons in aged mice compared to young (63% decrease) and middle-aged (62% decrease) mice (Figure 2I). Moreover, we identified significantly lower *Th*^-^/*Vglut2*^+^ cell density in the SNc compared to VTA. Aging also significantly impacted *Vglut2* mRNA expression within individual neurons. There were 49% fewer *Vglut2* mRNA grains per cell in aged mice compared to young animals and 17% fewer *Vglut2* mRNA grains per cell in middle-aged compared to young mice (Figure 2J). Finally, the absence of age-related decreases in DAPI-labeled cells (Figure 1I) strongly suggests continued resilience of midbrain glutamatergic neurons despite the decreases in *Vglut2* mRNA expression across time. Further, we show that the age-related decreases in *Vglut2* mRNA expression are not only limited to *Th*^+^/*Vglut2*^+^ DA neurons but occur more generally across the larger population of midbrain glutamatergic cells.

### Th and Vglut2 protein expression in projections to mouse striatum remain unchanged across aging

Since total numbers of ventral midbrain cells remained unchanged across all age groups, we examined whether dopaminergic and glutamatergic cells maintain Th and Vglut2 protein levels despite age-related decreases in mRNA expression. We specifically focused on the caudate/putamen (CPu) and nucleus accumbens (NAc), the primary projection sites of SNc and VTA dopaminergic neurons where Th and Vglut2 proteins are predominantly expressed in mice (Figure 3A-B).

**Figure 3.**
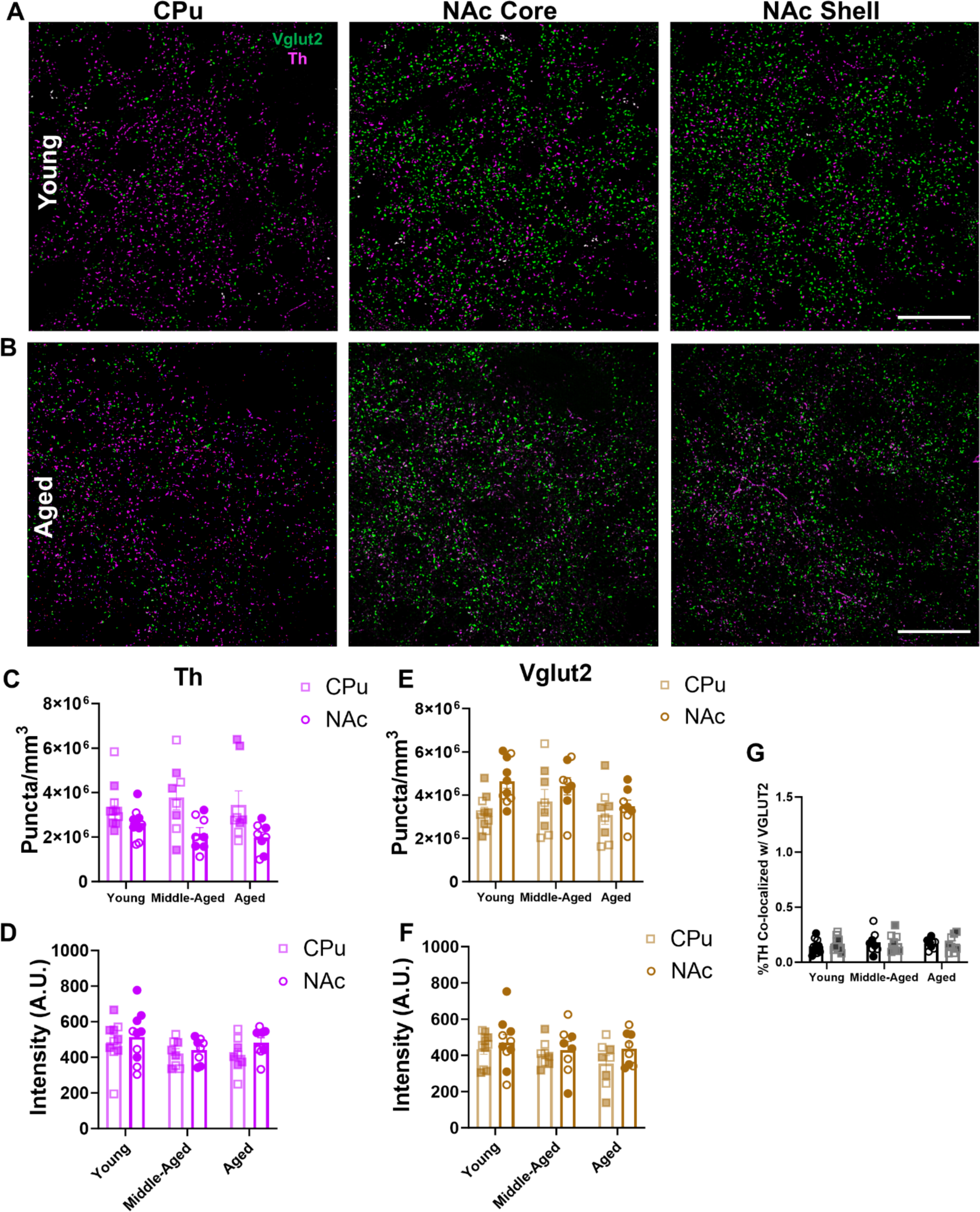
Th and Vglut2 protein expression is unaltered across aging in mouse striatum. **(A-B)** 60× representative images of immunohistochemical labeling of Th (in magenta) and Vglut2 (in green) protein in the CPu, NAc core, and NAc shell subregions of mouse striatum; scale bars=25μm. **(C)** There was no change in Th^+^ puncta density across aging. There were significantly higher levels of Th^+^ puncta in the CPu compared to the NAc (F_2,46_=0.3, P=0.74 for effect of age; F_1,46_=16.3, P=0.0002 for effect of region; F_2,46_=0.8, P=0.46 for effect of interaction). **(D)** There was no change in Th^+^ puncta intensity across aging or between striatal regions (F_2,46_=2.1, P=0.13 for effect of age, F_1,46_=1.7, P=0.20 for effect of brain region; F_2,46_=0.23, P=0.79 for effect of interaction). **(E)** There was no change in Vglut2^+^ puncta density across age. There were significantly higher levels of Vglut2^+^ puncta in the NAc compared to CPu (F_2,46_=2.2, P=0.12 for effect of age; F_1,46_=7.6, P=0.0083 for effect of region; F_2,46_=1.1, P=0.35 for effect of interaction). **(F)** There was no change in Vglut2^+^ puncta intensity across age or striatal region (F_2,46_=1.1, P=0.33 for effect of age; F_1,46_=2.2, P=0.14 for effect of region; F_2,46_=0.3, P=0.74 for effect of interaction). **(G)** There were low levels of TH co-localization with Vglut2 (∼0.2%) that remained unchanged with age or across NAc sub-region (F_2,46_=0.3, P=0.71 for effect of age; F_1,46_=0.1, P=0.79 for effect of NAc sub-region; F_2,46_=0.2, P=0.84 for effect of interaction). Shaded symbols represent female animals and open symbols represent males. Bars represent mean±SEM with points representing individual animals; N=8-10 per group.

Unlike the age-dependent decreases in *Th* mRNA, immunohistochemical labeling of Th protein demonstrated no change in Th^+^ puncta density in the striatum over the course of aging (Figure 3C). However, we did observe a difference in Th^+^ puncta density between regions, with higher levels of Th^+^ puncta in the CPu compared to the NAc. Moreover, we found no changes in average Th^+^ puncta intensity across age or between brain regions (Figure 3D). These findings suggest that Th protein expression in striatal projections is maintained across aging.

In glutamatergic projections to striatum, we similarly found no changes to Vglut2^+^ puncta density across aging (Figure 3E). We identified a significant difference between brain regions, with higher Vglut2^+^ puncta density in the NAc compared to the CPu. These data suggest denser NAc innervation from glutamatergic neurons, including Th^+^/Vglut2^+^ cells, consistent with recent reports^51^. Additionally, there were no differences between age groups or striatal regions in terms of average Vglut2^+^ puncta intensity (Figure 3F). This suggests that Vglut2 protein expression within striatal projections remained stable across aging.

Lastly, we also focused on Th and Vglut2 protein co-expression in striatal projections from midbrain Th^+^/Vglut2^+^ neurons which preferentially project to the medial NAc shell^47, 51–55^. We mainly found separate Th^+^ and Vglut2^+^ puncta along axonal projections within the mouse medial NAc shell, consistent with discrete sites of DA and glutamate release. Moreover, there were no differences in Th and Vglut2 protein co-localization across age groups or NAc sub-regions (*i.e.*, shell versus core) (Figure 3G). Together, our results indicate no change in either Th or Vglut2 protein expression in mouse striatum over the course of aging.

### *TH* and *VGLUT2* mRNA expression decrease with age in human midbrain

To translate our rodent findings to humans, we examined the impact of aging on *TH* and *VGLUT2* mRNA expression in VTA and SNc from postmortem human brain samples acquired from young, middle-aged, and aged subjects with no known lifetime psychiatric or neurological diseases (Table 1, Figure 4A-C). We first examined dopaminergic neurons in the midbrain. There was a significant effect of age group on the cell density of purely dopaminergic *TH*^+^/*VGLUT2*^-^ neurons (henceforth labeled as *TH*^+^) (Figure 4D), with aged subjects having 93% fewer *TH*^+^ neurons compared to young subjects. These data indicate substantial loss of *TH* mRNA expression as a function of age which recapitulated our mouse findings. Consistent with this, age and *TH*^+^ cell density showed a significant negative correlation (Supplementary Figure 7). There were also no observed differences in *TH* mRNA expression across brain region (Figure 4D), suggesting that the VTA and SNc were similarly affected throughout aging. Likewise, we found no changes across age group or brain region on *TH* mRNA grain levels within *TH*^+^ cells (Figure 4E).

**Figure 4.**
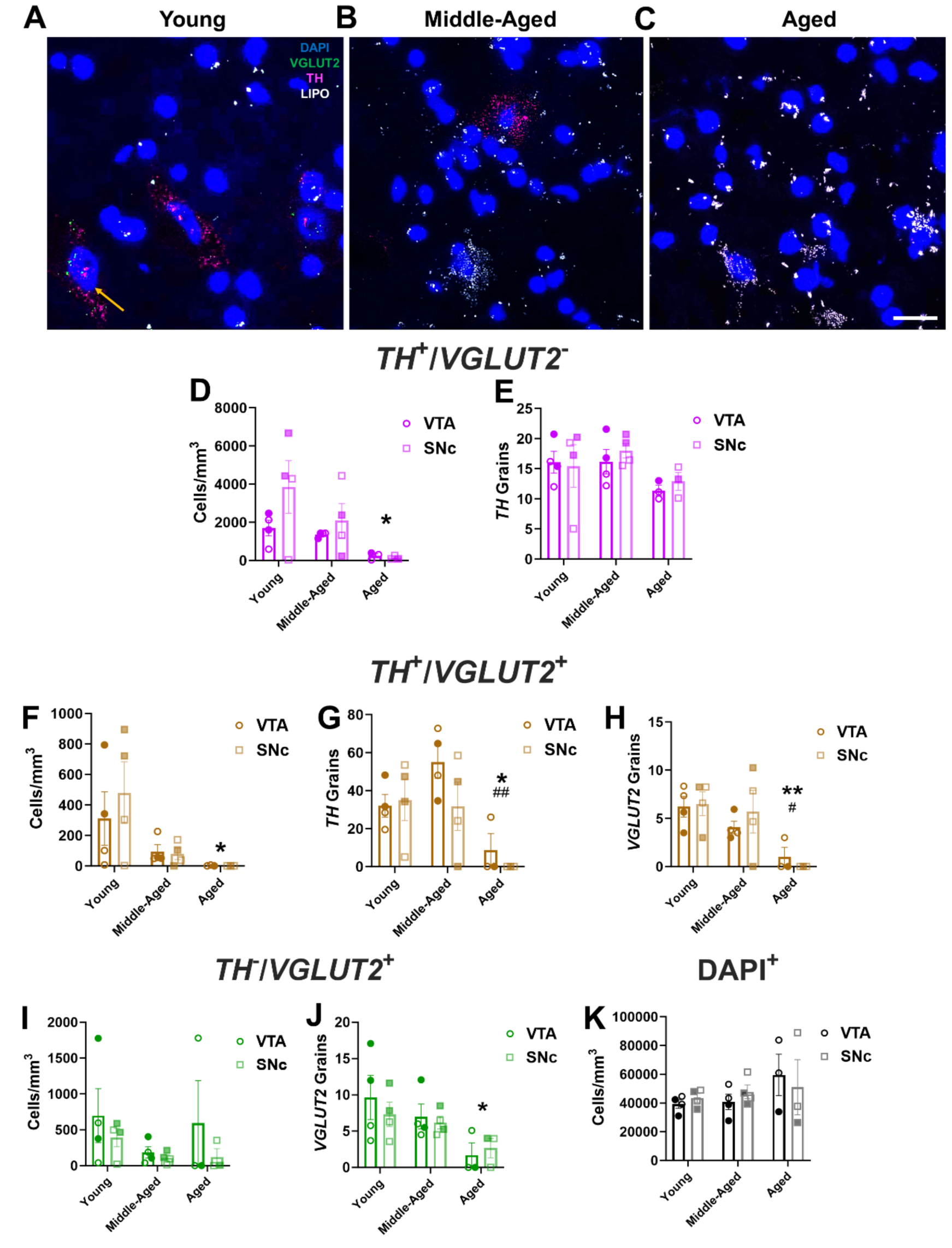
Age-related decreases in *TH* and *VGLUT2* mRNA expression in human VTA and SNc. **(A-C)** Representative 60× multiplex RNAscope images showing mRNA expression of TH (in magenta) and VGLUT2 (in green) mRNA expression alongside DAPI-labeled nuclei (in blue) in postmortem midbrains of young, middle-aged, and aged subjects; lipofuscin (LIPO, in white) was also present in middle-aged and aged tissue; scale bar=25μm. **(D)** There was an age-related decrease in cell density of non-glutamatergic *TH*^+^ neurons (*TH*^+^/*VGLUT2*^-^) (F_2,16_=5.2, P=0.018 for effect of age, F_1,15_=2.1, P=0.16 for effect of brain region; F_2,15_=1.1, P=0.37 for effect of interaction). **(E)** *TH* mRNA grain numbers in *TH*^+^ neurons did not change between age groups or across region (F_2,15_=2.5, P=0.12 for effect of age; F_1,15_=0.2, P=0.63 for effect of brain region; F_2,15_=0.2, P=0.83 for effect of interaction). **(F)** *TH*^+^/*VGLUT2*^+^ cell density diminished with age (F_2,16_=5.6, P=0.014 for effect of age; F_1,16_=0.2, P=0.63 for effect of region; F_2,16_=0.4, P=0.7 for effect of interaction). **(G)** *TH* mRNA grains per cell declined across aging (F_2,16_=8.6, P=0.0029 for effect of age; F_1,16_=1.6, P=0.23 for effect of region; F_2,16_=1.1, P=0.36 for effect of interaction). **(H)** *VGLUT2* mRNA grains per cell decreased across aging (F_2,16_=9.7, P=0.0018 for effect of age; F_1,16_=0.1, P=0.78 for effect of brain region; F_2,16_=0.5, P=0.63 for effect of interaction). **(I)** Among non-dopaminergic VGLUT2^+^ midbrain neurons (*TH*^-^/*VGLUT2*^+^), there was no significant change in cell density (F_2,16_=1.2, P=0.32 for effect of age; F_1,16_=1.7, P=0.21 for effect of region; F_2,16_=0.27, P=0.77 for effect of interaction). **(J)** Numbers of *VGLUT2* mRNA grains per cell were significantly decreased in aged subjects compared to young subjects (F_2,16_=5.1, P=0.019 for effect of age; F_1,16_=0.2121, P=0.65 for effect of region; F_2,16_=0.34, P=0.72 for effect of interaction). **(K)** There was no change in the density of DAPI-stained nuclei with age (F_2,16_=1.3, P=0.29 for effect of age; F_1,16_=0.02, P=0.90 for effect of region; F_2,16_=0.4, P=0.66 for effect of interaction). Shaded symbols represent female subjects and open symbols represent male subjects. Bars represent mean±SEM with points representing individual subjects; N=3-4 per group. *P<0.05 compared to young subjects. ^#^P<0.05 compared to middle-aged subjects.

We next examined effects of age on the cell density of *TH*^+^/*VGLUT2*^+^ coexpressing neurons. We discovered that there was a significant decrease in the density of *TH*^+^/*VGLUT2*^+^ mRNA coexpressing cells across age but no differences between brain region (Figure 4F). Aged subjects had 99% lower cell density of *TH*^+^/*VGLUT2*^+^ neurons compared to the young group, which mirrors our findings in mice. This was similarly coupled with a negative correlation between age and *TH*^+^/*VGLUT2*^+^ cell density in the SNc (Supplementary Figure 7). For both *TH* and *VGLUT2* mRNA grain levels within co-expressing cells, there was a significant decrease with age but no effect of brain region (Figures 4G, 4H). Aged subjects had 90% fewer *TH* mRNA grains (Figure 4G) and 90% fewer *VGLUT2* grains (Figure 4H) in coexpressing cells compared to middle-aged subjects. Moreover, aged subjects also had 87% fewer *TH* grains (Figure 4G) and 92% fewer *VGLUT2* grains (Figure 4H) versus young subjects.

We additionally examined the impact of aging on purely glutamatergic *TH*^-^/*VGLUT2*^+^ midbrain neurons in our human subjects. In contrast to dopaminergic cells, there were no differences across age or brain region on *TH*^-^/*VGLUT2*^+^ neuron density (Figure 4I). On the other hand, there was an effect of age on *VGLUT2* mRNA expression, with a 74% decrease in *VGLUT2* mRNA grains per cell in aged versus young subjects (Figure 4J).

To determine whether age-related decreases in *TH*^+^ and *VGLUT2*^+^ cell density were associated with midbrain DA neuron loss, we quantified DAPI^+^ nucleus density in human midbrain across aging. As in mice, we found no changes between age group or region on DAPI-stained nuclear density in human midbrain (Figure 4K). This indicated that cells continue to survive in human midbrain throughout aging despite significant reductions in the expression of *TH* and/or *VGLUT2* mRNA. Just as importantly, our findings appear to be strongly conserved across species (Figure 1I).

### Impact of aging on midbrain projections to the human striatum

As in mice, we investigated whether midbrain neuronal TH and VGLUT2 protein expression was preserved across aging. We therefore measured TH and VGLUT2 protein expression in midbrain neuron projections to the CPu and NAc of the same young, middle-aged, and aged human subjects (Figure 5A-C). In contrast to mice, there was a decrease in total TH^+^ puncta density in humans (Figure 5D), albeit to a lesser degree compared to the age-related diminishment of *TH* mRNA expression (Figure 4). Indeed, there was a significant 30% decrease in TH^+^ puncta density in aged subjects compared to middle-aged subjects as well as a trend (25% decrease) in aged versus young subjects (P=0.068). Moreover, consistent with our mouse data, there was no change in TH^+^ puncta intensity across age or region (Figure 5E). In the context of lower TH^+^ puncta density in aged striatum, conservation of TH^+^ puncta intensity suggests that the TH protein expression in the remaining terminals remains either the same or becomes elevated. There may therefore be compensatory mechanisms that maintain TH protein levels in remaining dopaminergic boutons of aged subjects.

**Figure 5.**
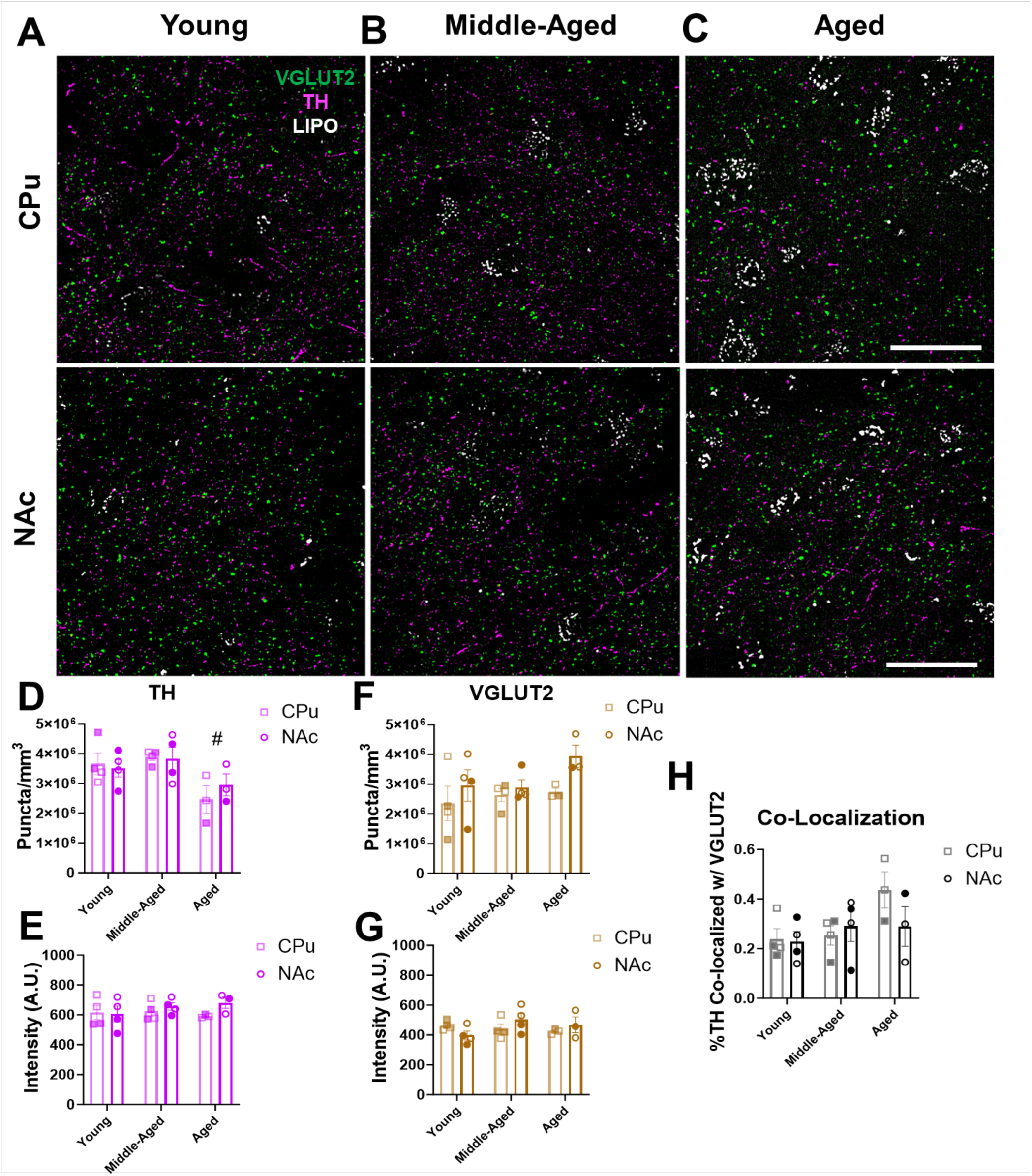
TH and VGLUT2 protein expression in projections to human striatum remain unchanged throughout aging. **(A-C)** 60× representative immunohistochemistry images of TH (in magenta) and VGLUT2 (in green) protein expression within the NAc and CPu subregions of human striatum; lipofuscin (in white) was also present in the images; scale bars=25μm. **(D)** There was an age-related decrease in the density of TH^+^ puncta in aged versus middle-aged subjects (F_2,16_=5.8, P=0.013 for effect of age; F_1,16_=0.1, P=0.74 for effect of region; F_2,16_=0.5, P=0.63 for effect of interaction). **(E)** There were no significant age-related changes in the intensity of TH^+^ puncta in the striatum across age or between regions (F_2,16_=0.3, P=0.72 for effect of age; F_1,16_=1.3, P=0.27 for effect of region; F_2,16_=0.8, P=0.47 for effect of interaction). **(F)** VGLUT2 puncta density remained unaltered across aging and between regions (F_2,16_=1.5, P=0.26 for effect of age; F_1,16_=4.2, P=0.057 for effect of region; F_2,16_=2.3, P=0.14 for effect of interaction). **(G)** VGLUT2 puncta intensity remained unaltered across aging and between regions (F_2,16_=0.8, P=0.46 for effect of age; F_1,16_=0.2, P=0.63 for effect of region; F_2,16_=2.3, P=0.14 for effect of interaction). **(H)** Quantification showed low levels of TH and VGLUT2 co-localization in projections to the striatum (∼0.2-0.4%) that did not change with age or across region (F_2,16_=2.7, P=0.097 for effect of age; F_1,16_=0.8, P=0.39 for effect of region; F_2,16_=1.4, P=0.28 for effect of interaction). Shaded symbols represent female subjects and open symbols represent male subjects. Bars represent mean±SEM with points representing individual subjects; N=3-4 per group. ^#^P<0.05 compared to middle-aged subjects.

We next analyzed the impact of aging on VGLUT2 protein expression in glutamatergic projections to human striatum. In contrast to dopaminergic projections, analysis of VGLUT2^+^ striatal puncta density revealed no changes across age groups or brain regions (Figure 5F). Similarly, there were no changes across age groups or brain regions in VGLUT2 intensity (Figure 5G). These findings suggest that both the total number of glutamatergic projections and VGLUT2 protein expression remain unchanged in the human striatum throughout normal aging. Finally, analysis of co-localization of TH^+^ and VGLUT2^+^ puncta in the CPu and NAc revealed ∼0.2-0.4% co-localization in young, middle-aged, and aged subjects (Figure 5H) which is consistent with our findings in mice (Figure 3H). These data are also in line with prior evidence of segregation of TH^+^ and VGLUT2^+^ vesicles in axonal boutons within the mammalian striatum^42, 56–58^. Furthermore, we found no differences in TH and VGLUT2 co-localization across age or between brain regions (Figure 5H). This indicates a lack of age-related changes in TH/VGLUT2 co-expression in axonal boutons.

### Ribosomal gene expression is maintained throughout aging in mouse midbrain

We posited that a translational response may be responsible for the maintenance of TH and VGLUT2 protein expression in human and mouse midbrain despite age-related decreases in mRNA expression. To test this, we initially focused on *Rpl6* which encodes the 60S ribosomal protein L6 (RPL6), a key component of the 60S large ribosomal subunit^59^. Significantly, in addition to its functions in protein translation, RPL6 plays critical roles in DNA damage repair^60^. Given the progressive accumulation of oxidative DNA damage during aging, RPL6’s ability to mitigate this damage is important for maintaining cellular resilience^60^. We first measured *Rpl6* mRNA expression across mouse midbrain throughout aging via multiplex RNAscope (Figures 6A, 6B). Unlike *Th* and *Vglut2* mRNAs, there was no net change in *Rpl6* mRNA grain numbers per cell across aging. We then compared average *Rpl6* expression according to cell type and midbrain region in young versus aged mice, focusing on midbrain *Th*^+^/*Vglut2*^-^, *Th*^+^/*Vglut2*^+^ and *Th*^-^/*Vglut2*^+^ cells (Figure 6C). We found no significant differences across age group or brain region. Finally, we quantified the density of *Rpl6* mRNA grains in both the VTA and SNc, and observed no significant differences across age or brain region (Figure 6D). Overall, in contrast to *Th* or *Vglut2* mRNA expression, these results show that *Rpl6* expression and density is preserved during the aging process, suggesting that Rpl6 may play a role in maintaining protein expression in midbrain neurons.

**Figure 6.**
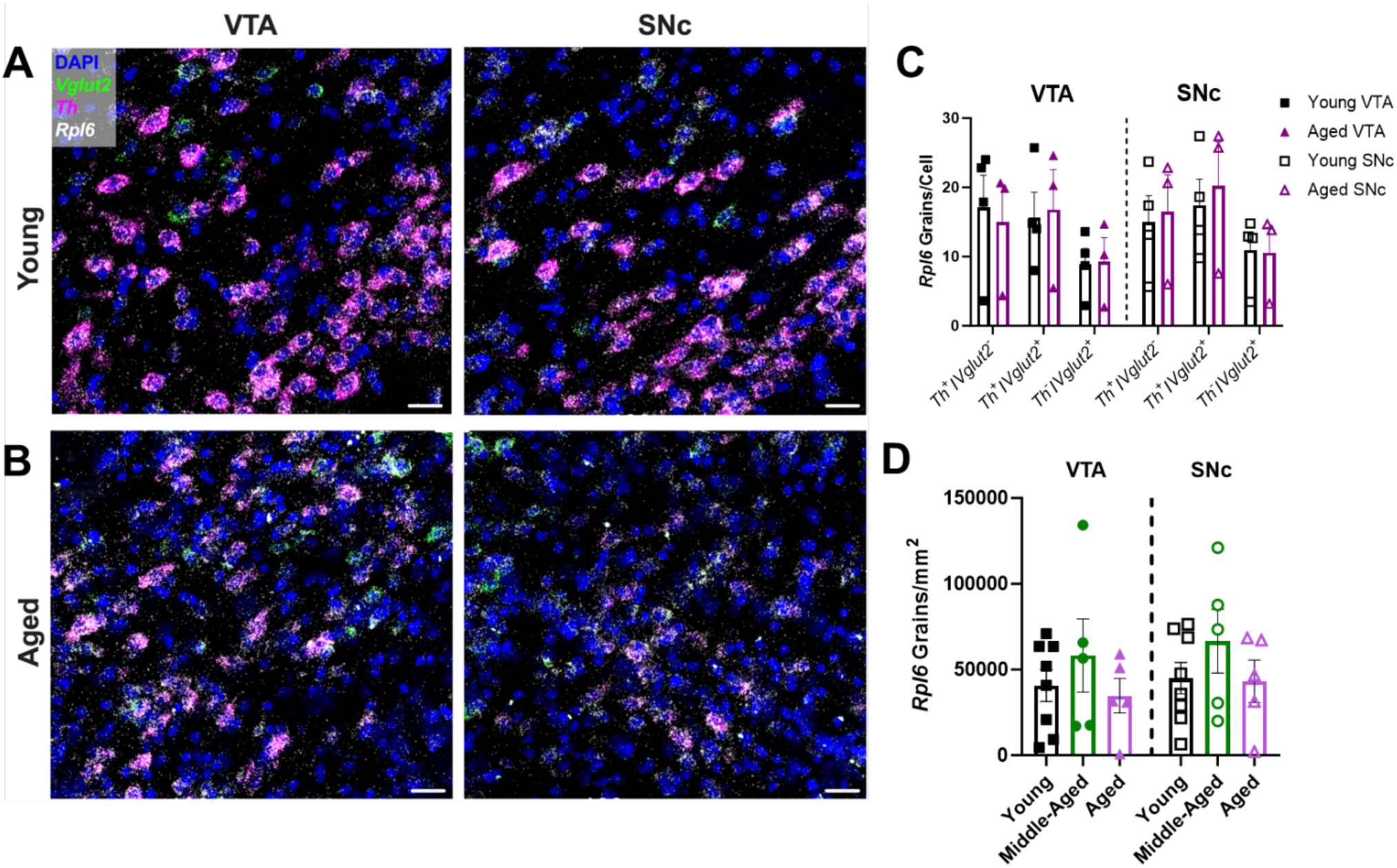
*Rpl6* mRNA expression is preserved across aging in the mouse midbrain. **(A-B)** Representative 60× multiplex RNAscope images of *Th* (in magenta), *Vglut2* (in green), and *Rpl6* (in white) mRNA expression alongside DAPI nuclear stain (in blue) in the VTA and SNc of young and aged wild-type mice. Scale bars=25μm **(C)** Cell type- and region-specific analysis of average *Rpl6* mRNA grain numbers per cell. There were no significant expression differences in *Th*^+^/*Vglut2*^-^, *Th*^+^/*Vglut*2^+^, and *Th*^-^/*Vglut2*^+^ cell types in either the VTA or SNc of young versus aged mice (F_1,30_=0.30, P=0.59 for effect of age; F_1,30_=0.05, P=0.82 for effect of region; F_2,30_=3.558, P=0.04 for effect of cell type; F_1,30_=0.10, P=0.75 for effect of age×region; F_2,30_=0.12, P=0.89 for effect of age×cell type; F_2,30_=0.09, P=0.92 for effect of region×cell type; F_2,30_=0.07, P=0.94 for effect of age×region×cell type. **(D)** Quantification of *Rpl6* mRNA density across both VTA and SNc showed no significant differences between the age groups (F_2,30_=1.6091, P=0.22 for effect of age; F_1,30_=0.4163, P=0.52 for effect of region; F_2,30_=0.018, P=0.98 for effect of interaction). Bars represent mean±SEM with points representing individual animals; N=3-8 per group.

We also explored the possibility that age-related maintenance of ribosomal gene expression is not limited to *Rpl6*, but instead may be more generalized throughout the family of ribosome-related genes. We posited that maintaining or upregulating ribosomal genes may represent a homeostatic response to diminished mRNA levels through aging by maintaining ribosomal translation. To address this, we performed transcriptomic analyses on previously deposited scRNA-seq data from brains of young versus aged wild-type mice^45^, focusing on *Th^+^* neurons. We did not additionally examine smaller subpopulations of DA neurons including *Th*^+^/*Vglut2*^+^ neurons since this dataset did not possess a sufficient number of these cells to power such subanalyses. Consistent with our *Rpl6* findings, there was no significant difference in overall ribosomal gene expression between young and aged mice (Figure 7). Of the 141 ribosomal genes analyzed, expression of the majority of genes was either maintained or even increased in aged *Th^+^* neurons (Supplemental Figure 8). Taken together, our data strongly suggest that expression of ribosome-related genes is broadly preserved across normal aging.

**Figure 7.**
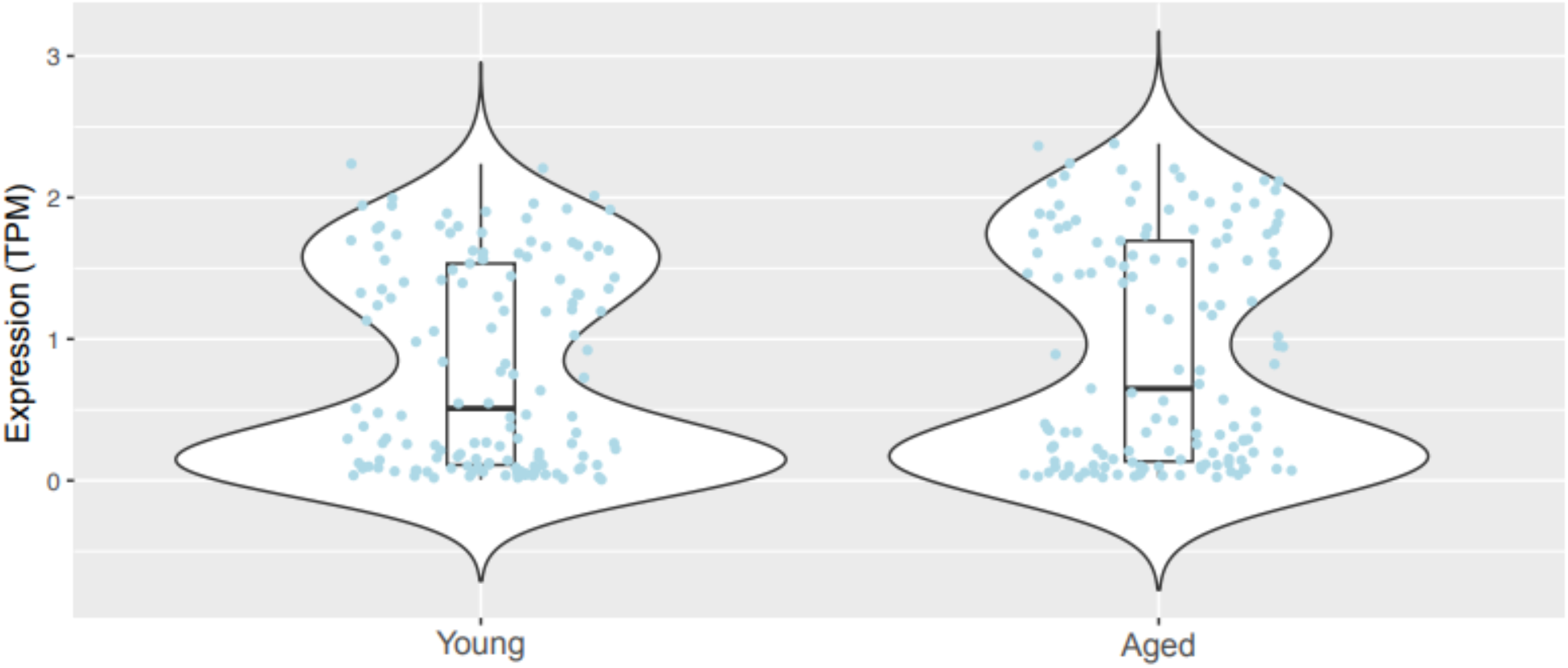
Transcriptomic analysis of ribosomal genes in young and aged mice. Expression levels of 141 ribosomal genes in *Th*^+^ cells (young cells, N=295; aged cells, N=502). Each jitter represents averaged expression levels (in TPM) for every assayed ribosomal gene in *Th*^+^ neurons of young versus aged wild-type mice. Each of the 141 ribosomal genes was expressed in at least 10 *Th*^+^ cells. There was no significant difference in gene expression between the young and old groups (t = 0.893, P = 0.37). Box and whiskers represent the 25^th^ percentile, median, and 75^th^ percentile.

## Discussion

The present study identified alterations in the coordination between mRNA and protein expression within midbrain DA and glutamate systems across aging in mice and humans. Taking advantage of multiplex RNAscope’s single-cell resolution, we showed an age-related decrease in *TH* mRNA expression in midbrain DA neurons. Yet, despite substantial age-related losses in mRNA expression in the VTA and SNc, we did not find accompanying decreases in TH protein expression or dopaminergic cell loss. Though earlier work linked diminished DA neuron pigmentation or TH expression during aging to neurodegeneration^17, 61, 62^, other studies used Nissl staining to show that the number of cell nuclei in the human and primate SNc are maintained with age^63, 64^. Our findings showing absence of age-related cell loss also contrast with PD models in mice or clinical PD which demonstrate stark decreases in total nuclei in both the VTA and SNc^65–68^.

In mice and humans, we also found that age-related decreases in mRNA expression were not limited to *TH* expression. Rather, there was significant loss of *VGLUT2* mRNA expression in midbrain *TH*^+^/*VGLUT2*^+^ DA neurons as well as in purely glutamatergic cells. Nevertheless, like TH in DA neurons, there were no significant age-related changes to VGLUT2 protein expression. These data strongly suggest compensatory mechanisms in response to altered mRNA expression that occur during normal aging. We posit that these mechanisms are likely part of a more global response employed by multiple midbrain neuronal populations. Yet, while there is a robust literature examining age-related changes to glutamatergic neurotransmission in neurodegenerative diseases like PD and Alzheimer’s disease^69, 70^, there are relatively few studies examining effects of normal aging on the glutamate system either in rodent models or humans. In rodents, there has been a general lack of consensus on effects of age on glutamate content across brain regions (*i.e.*, decreases, no change, or even increases over time)^69, 71, 72^. However, both *in vitro* and *in vivo* studies have consistently demonstrated no significant age-related alterations in basal and stimulated glutamate release including in cortex and striatum^69, 73, 74^. Such findings are consistent with our results demonstrating no age-related changes in VGLUT2’s protein expression, enabling stable glutamate neurotransmission throughout aging.

Our mapping of mouse and human TH^+^/VGLUT2^+^ DA/glutamate neuron distribution in the midbrain demonstrates that the anatomic distribution of these neurons is conserved across species^48, 49, 75–77^. Just as importantly, this work provides one of the first comprehensive maps of TH^+^/VGLUT2^+^ neurons using both protein and mRNA expression within human midbrains of males and females. These results are in line with previous findings in rodents, non-human primates, and humans showing that VGLUT2^+^ DA neurons are preferentially localized to the medial VTA^47–49, 58^. Our mapping data also shed further light on the ongoing debate whether DA and glutamate are released from the same axonal sites. Though we found both TH and VGLUT2 protein localized to the same axons, we found little direct co-localization at the same sites. These data suggest that DA and glutamate may be released from different sites within the axons of these neurons, consistent with previous findings^42, 56, 57, 78, 79^. However, we cannot rule out that additional projections from TH^+^/VGLUT2^+^ cells which project to extra-striatal sites (*e.g*., frontal cortex and hippocampus)^47, 52, 80^ may have greater co-localization of TH and VGLUT2.

What are the biological mechanisms that contribute to loss of *TH* and *VGLUT2* mRNA expression and the associated compensatory responses during aging? As organisms age, the steady accumulation of extrinsic environmental and intrinsic insults leads to elevated cell stress and gradual impairments in cell function^81^. For example, repeated exposures to cytotoxic reactive oxygen species (ROS) throughout aging causes DNA damage. This leads to a gradual degradation of genomic integrity which, in neurons, may eventually lead to neurodegenerative disease pathology^82, 83^. DA neurons are especially susceptible to oxidative stress given the conversion of DA into intermediates that further raise ROS levels, contributing to PD pathogenesis^84, 85^. Cells may also respond to insults during aging by changing the epigenetic landscape to alter gene expression^86^. We propose that a combination of these age-related genomic and epigenomic modifications may contribute to diminished efficiency of gene transcription of both *TH* and *VGLUT2* that result in their lower mRNA expression. On the other hand, our findings also point to an age-related homeostatic process that maintains protein expression despite decreases in mRNA expression, enabling neurons to maintain the relative stability of midbrain neurotransmission. Specifically, we examined potential ribosomal contributions to the preservation of protein expression despite diminished mRNA levels in dopaminergic and glutamatergic neurons. Our analysis of a comprehensive scRNA-seq dataset derived from young and aged mice^45^ confirmed the existance of such a ribosomal mechanism by demonstrating that expression of ribosomal genes was broadly maintained or even upregulated in aged dopaminergic neurons. In contrast, the same dataset showed downregulation of many non-ribosomal genes including *Th*, consistent with our data. In contrast, age-related neurodegenerative disorders like PD result in downregulation of select ribosomal genes. RPL6, a key component of the ribosomal translational machinery^59, 87^, is downregulated in PD^87^, but our data showed that *Rpl6* expression is maintained across the course of healthy aging. This raises the possibility that the pathology associated with age-related neurodegenerative disorders is driven and/or exacerbated by a diminished capacity of the translational machinery to keep up with declines in mRNA expression during aging. In humans, transcriptional profiling also demonstrated age-associated increases in expression of ribosomal genes in muscle^88^. This points to an age-related ribosomal mechanism to preserve the integrity of the cellular proteome that may be more generalized throughout the body. Preservation of ribosomal gene expression during aging is also evident in flies, further emphasizing that these mechanisms are likely strongly conserved across evolution. Indeed, the *Drosophila* model demonstrated broadly diminished expression of most genes in the adult brain during aging unlike ribosomal genes which remained protected from similar declines^89^.

How are ribosomal genes protected from the age-related declines in expression observed for *TH* and *VGLUT2*? Unlike most genes, ribosomal genes are under the control of a unique core promoter that employs the TCT motif which encompasses the transcriptional start site and is therefore required for ribosomal gene transcription^90^. This TCT-based transcriptional system is conserved from flies to mammals, and has been suggested to play a key role in explaining why ribosomal gene expression does not drop during aging^89, 90^. Further work is required to elucidate the mechanisms at the transcriptional level responsible for this age-associated selective preservation of ribosomal gene expression.

The balance between protein degradation and protein synthesis may offer an additional mechanism to maintain stable protein levels during healthy aging. Indeed, proteasome-mediated proteolytic activity diminishes with age^91–95^. By decreasing protein degradation relative to rates of *de novo* protein synthesis, TH and VGLUT2 protein can persist longer in aged neurons to forestall potential alterations in neurotransmission. Disrupting the proteostatic balance can therefore precipitate development of abnormal protein aggregation (*e.g.*, α-synuclein aggregates) that gradually contributes to eventual neurodegenerative pathologies^96^.

In conclusion, this study shows loss of coordination between mRNA and protein expression during aging that is highly conserved between mouse and human midbrain. Significantly, our results also suggest age-associated homeostatic mechanisms that maintain protein expression by maintaining expression of ribosomal genes. Ultimately, these findings highlight an important set of biological changes associated with aging within both mouse and human DA and glutamate systems. Therefore, developing a deeper understanding of these aging phenomena may pave the way for new, more effective therapeutic approaches to boost neuron resilience and prevent neurodegeneration.

## Acknowledgements

We are grateful for discussions and technical assistance provided by Emma O’Leary, Tyler Fortuna, and Eric Zimmerman. This study was supported by The Pittsburgh Foundation (John F. and Nancy A. Emmerling Fund of the Pittsburgh Foundation, FPG00043 to Z.F.), the Commonwealth of Pennsylvania (PA-HEALTH to Z.F.), and the National Institutes of Health (R21AG068607 to Z.F.; R01ES034037 to Z.F.; F31NS11811 to S.B., R01ESR36DA057972 to J.G., T32GM133353 to C.W. and J.G.K., T32MH019986 to S.J.M.).

## Author Contributions

S.A.B. and Z.F. conceived the project. S.A.B., S.J.M., J.R.G., C.W., T.B.T., J.G.K., K.N.F., and R.W.L. performed imaging experiments and data analysis. J.R.G. and D.A.L. prepared postmortem human brain samples. C.F. and R.W.L. performed the transcriptomic analyses. S.A.B., S.J.M., J.G.K., and Z.F. wrote the manuscript with contributions from co-authors.

