## Supplementary Materials for "Aging disrupts the coordination between mRNA and protein expression in mouse and human midbrain"

Includes Supplementary Figures 1-8

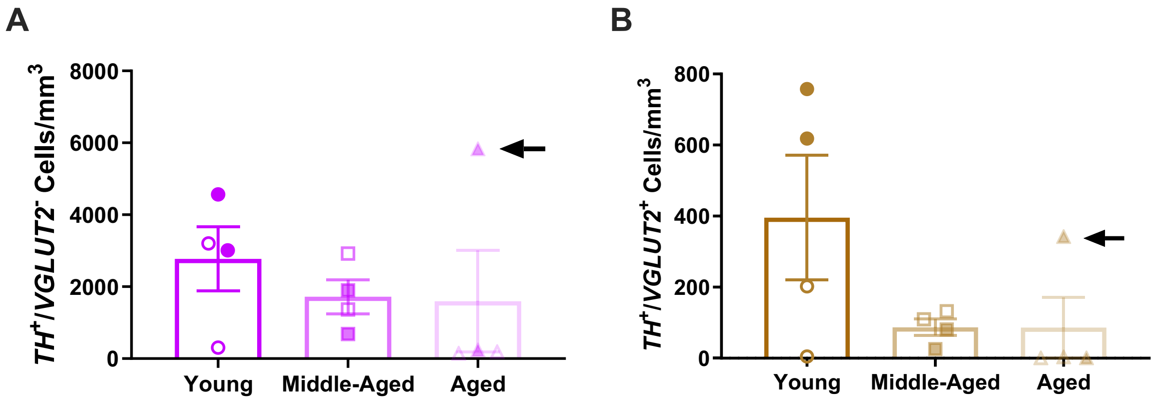


**Supplementary Figure 1. Human midbrain *TH* and *VGLUT2* mRNA expression including outlier. (A)** *TH*$^{+}$/*VGLUT2*$^{-}$ and **(B)** *TH*$^{+}$/*VGLUT2*$^{+}$ cell densities in young, middle-aged, and aged postmortem human ventral midbrain, showing the outlier aged subject. Shaded symbols represent female subjects and open symbols represent male subjects. Outlier is denoted by the black arrow. Bars represent mean±SEM with points representing individual subjects.

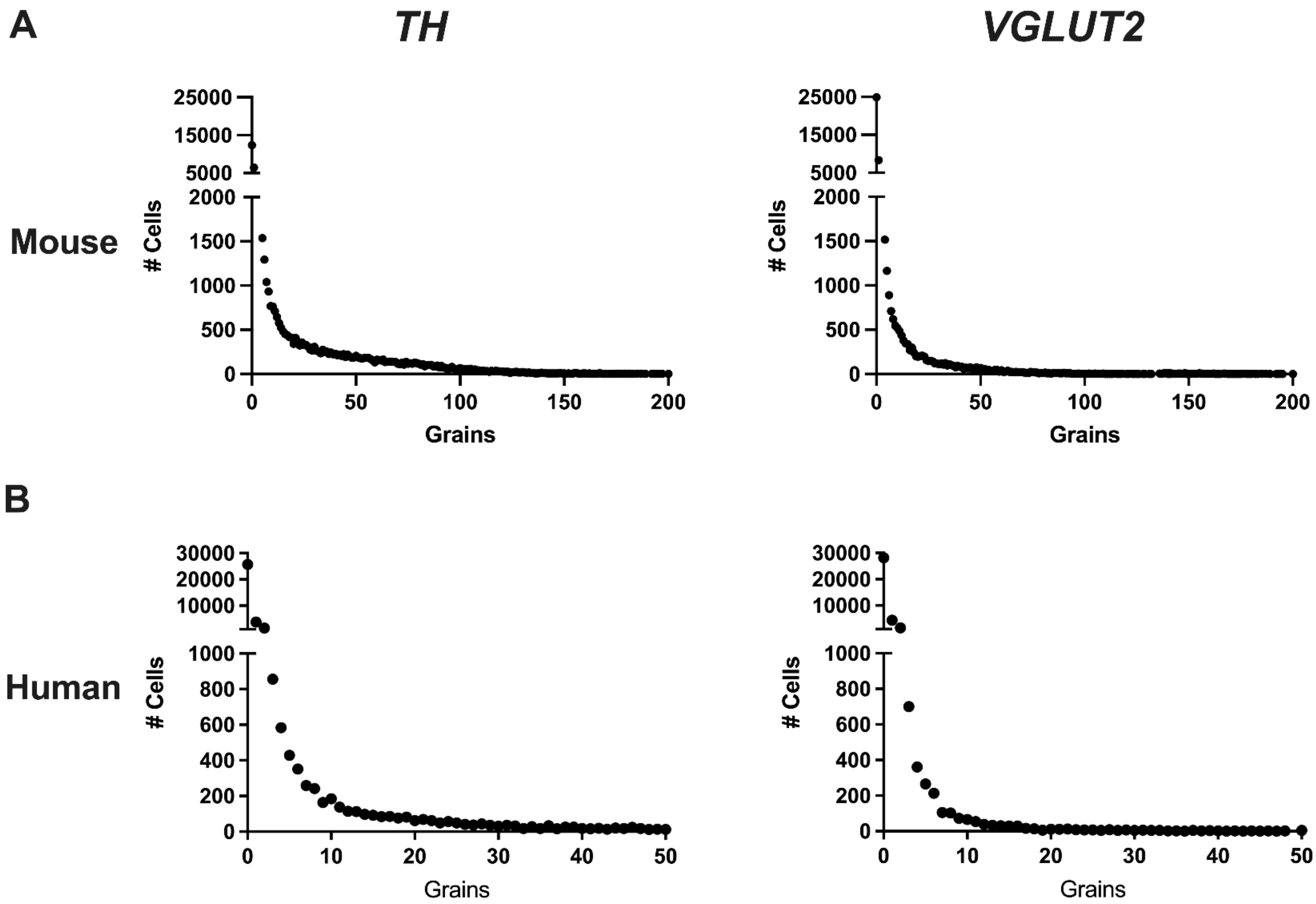


**Supplementary Figure 2. Histograms of *TH* and *VGLUT2* mRNA expression levels.** Histograms of *TH* and *VGLUT2* mRNA expression levels of all cells analyzed in **(A)** mice and **(B)** human subjects.

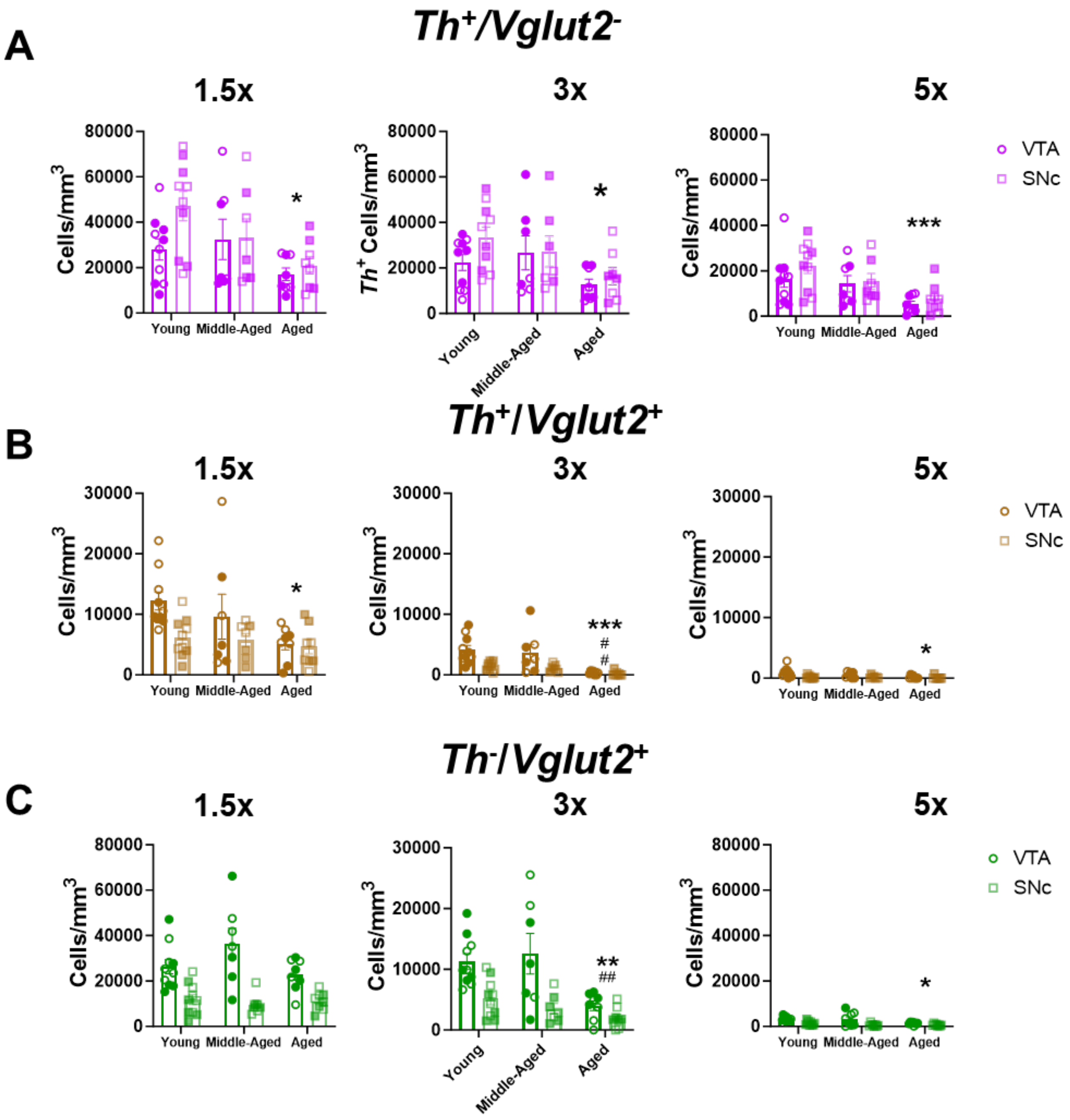


**Supplementary Figure 3. Positive cell threshold determination for mRNA expression analyses in mouse midbrain according to cell type.** Positive cell determination thresholds of 1.5x, 3x and 5x above background levels for **(A)** *Th*⁺/*Vglut2*⁻, **(B)** *Th*⁺/*Vglut2*⁺, and **(C)** *Th*⁻/*Vglut2*⁺ cell densities. Shaded symbols represent female subjects and open symbols represent male subjects. Bars represent mean±SEM with points representing individual animals; N=7-10 per group. *$P<0.05$, **$P<0.01$ ***$P<0.001$ compared to young, ##$P<0.01$ compared to middle-age.

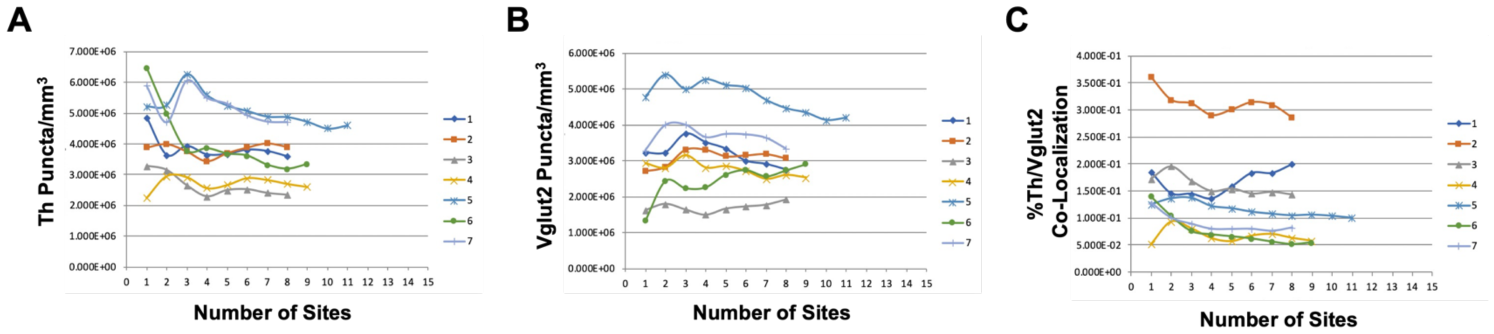


**Supplementary Figure 4. Running means analysis of a random sample of mouse striatal slices.** Analysis of Th and Vglut2 immunohistochemical labeling in the mouse striatum from 7 samples. These data showed that the means of $Th^+$ puncta density **(A)**, $Vglut2^+$ puncta density **(B)**, and percentage of $Th^+$ puncta that co-localized with Vglut2 **(C)** stabilized once 8 random sites had been quantified. This enabled acquisition of a reliable mean of the dependent measures in the assayed striatal area.

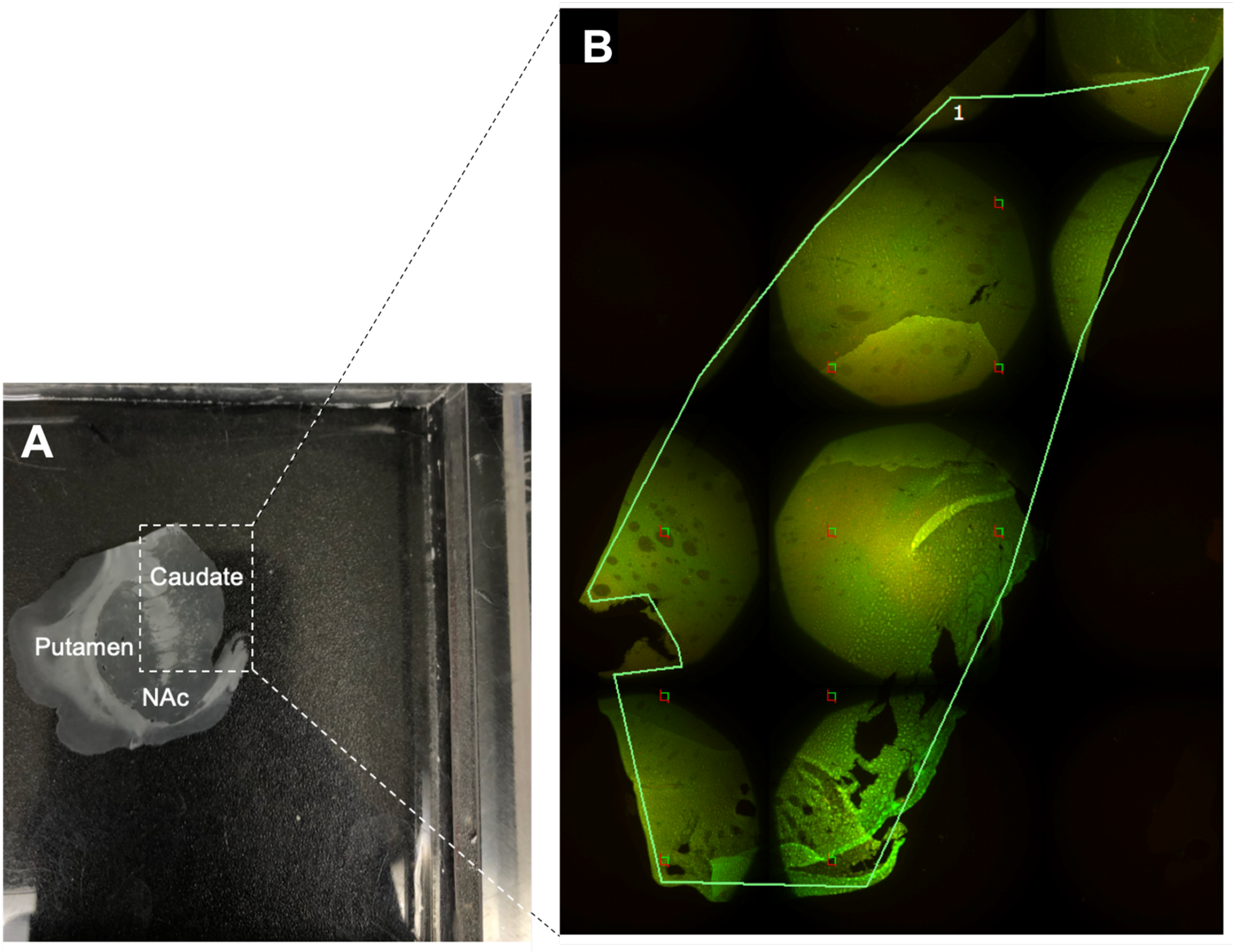


**Supplementary Figure 5. Example of human striatal section sampling for immunohistochemical imaging. (A)** Representative image of a caudate slice from human striatum used for immunohistochemistry studies. **(B)** 4x montage image of the same human striatal slice with the region of interest drawn and stereological selection of 100 μm × 100 μm sampling boxes separated by 2000 μm shown in the green and red squares. The areas within each sampling box were imaged and analyzed for TH and VGLUT2 protein expression.

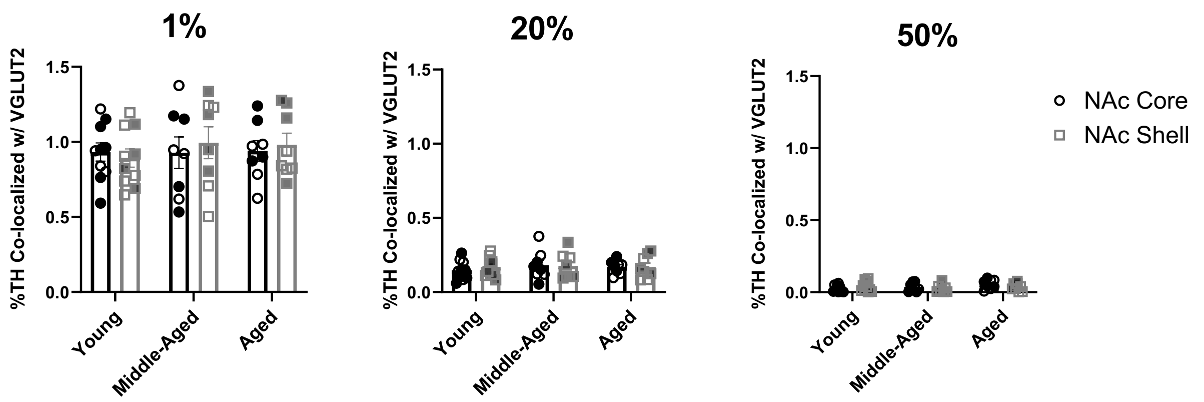


**Supplementary Figure 6. TH co-localization with VGLUT2 at different overlap thresholds in mice.** There were no significant differences between age groups and brain regions for any threshold (P>0.05). Shaded symbols represent female subjects and open symbols represent male subjects. Bars represent mean±SEM with points representing individual animals.

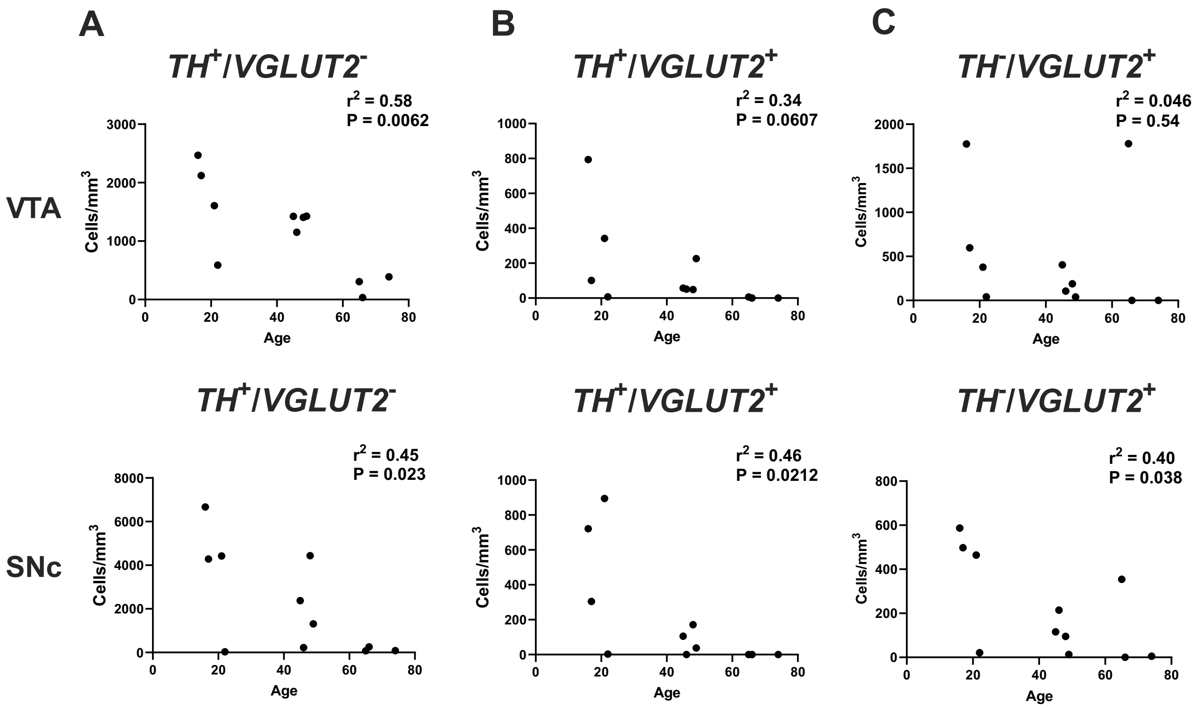


**Supplemental Figure 7. Relationships between cell type density and age in human midbrain.** Correlations were determined between age and cell densities for the following midbrain cell types: **(A)** *$TH^+$/$VGLUT2^-$* (VTA: $r^2$=0.58, P=0.006; SNc: $r^2$=0.45, P=0.02) **(B)** *$TH^+$/$VGLUT2^+$* (VTA: $r^2$=0.34, P=0.06; SNc: $r^2$=0.46, P=0.02) **(C)** *$TH^-$/$VGLUT2^+$* (VTA: $r^2$=0.046, P=0.54; SNc: $r^2$=0.40, P=0.038) in the VTA and SNc across all the human subjects.

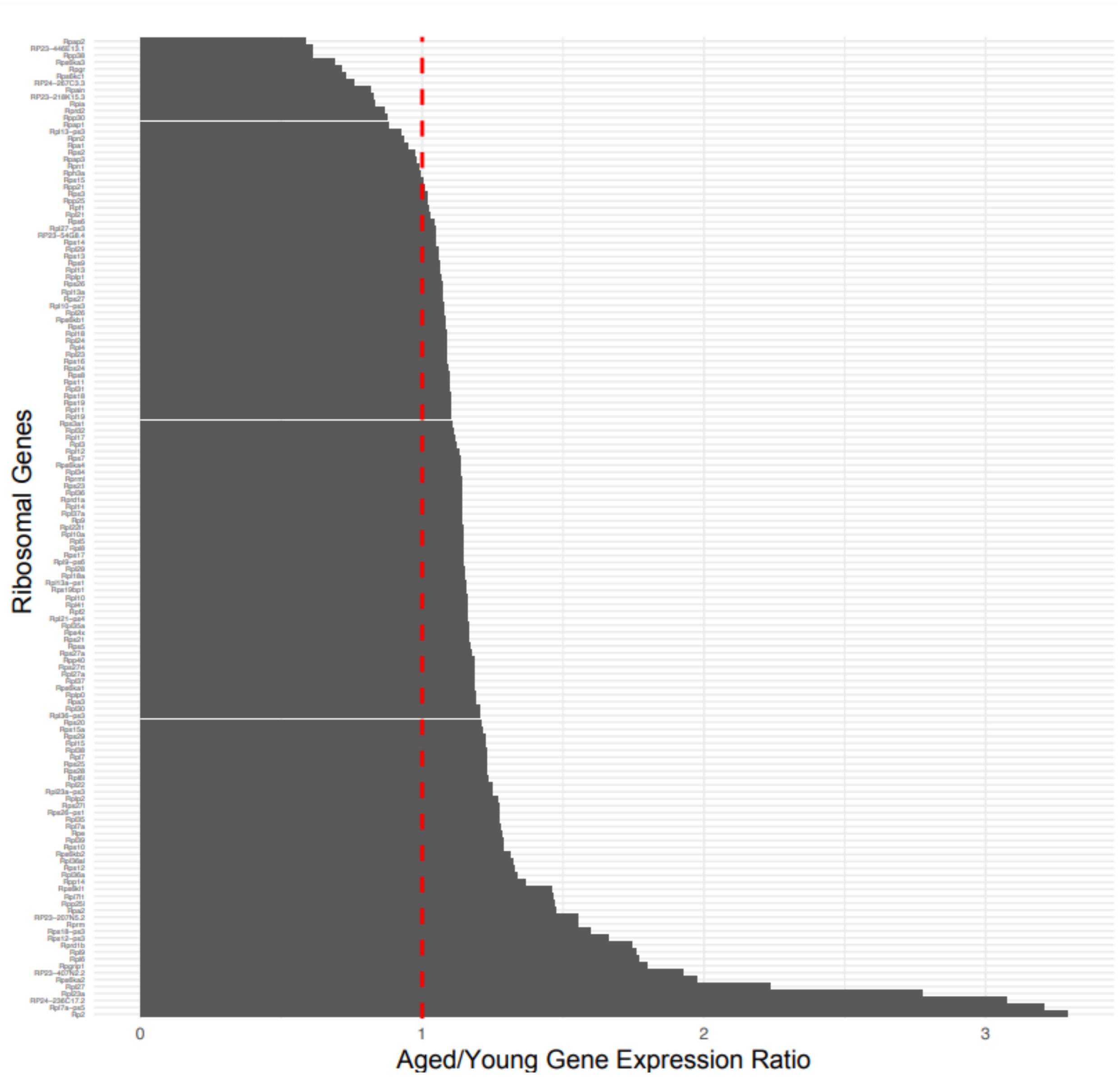


**Supplemental Figure 8. Aged/young gene expression ratios for ribosomal genes in *Th*⁺ neurons.** All 141 ribosomal gene expression ratios (Aged/Young) in *Th*⁺ cells were graphed as bar plots. Each bar represents individual ribosomal genes. The red dashed line indicates equal expression in aged and young groups. Every ribosomal gene was expressed in N≥10 *Th*⁺ cells.
